# Predicting Functional Effects of Missense Variants in Voltage-Gated Sodium and Calcium Channels

**DOI:** 10.1101/671453

**Authors:** Henrike O. Heyne, David Baez-Nieto, Sumaiya Iqbal, Duncan Palmer, Andreas Brunklaus, the Epi25 Collaborative, Katrine M. Johannesen, Stephan Lauxmann, Johannes R. Lemke, Rikke S. Møller, Eduardo Pérez-Palma, Ute Scholl, Steffen Syrbe, Holger Lerche, Patrick May, Dennis Lal, Arthur J. Campbell, Jen Pan, Hao-Ran Wang, Mark J. Daly

## Abstract

Malfunctions of voltage-gated sodium and calcium channels (SCN and CACNA1 genes) have been associated with severe neurologic, psychiatric, cardiac and other diseases. Altered channel activity is frequently grouped into gain or loss of ion channel function (GOF or LOF, respectively) which is not only corresponding to clinical disease manifestations, but also to differences in drug response. Experimental studies of channel function are therefore important, but laborious and usually focus only on a few variants at a time. Based on known gene-disease-mechanisms, we here infer LOF (518 variants) and GOF (309 variants) of likely pathogenic variants from disease phenotypes of variant carriers. We show regional clustering of inferred GOF and LOF variants, respectively, across the alignment of the entire gene family, suggesting shared pathomechanisms in the SCN/CACNA1 genes. By training a machine learning model on sequence- and structure-based features we predict LOF- or GOF-associated disease phenotypes (ROC = 0.85) of likely pathogenic missense variants. We then successfully validate the GOF versus LOF prediction on 87 functionally tested variants in *SCN1/2/8A* and *CACNA1I* (ROC = 0.73) and in exome-wide data from > 100.000 cases and controls. Ultimately, functional prediction of missense variants in clinically relevant genes will facilitate precision medicine in clinical practice.

## Introduction

Voltage-gated sodium (Na_v_s) and calcium channels (Ca_v_s) play a critical role in initiating and propagating action potentials across a broad variety of excitable cells and physiological functions. Upon membrane depolarisation, Na_v_s and Ca_v_s are activated and inactivated within milliseconds, leading to a transient influx of sodium or calcium ions into the cell (Catterall, 1995). Genes encoding the channel protein (in humans, Na_v_s’ channel proteins are encoded by 10 SCN and Ca_v_s’ channel proteins are encoded by 10 CACNA1 genes) have been associated with multiple predominantly neurological and neurodevelopmental diseases. These diseases include developmental and epileptic encephalopathy (*SCN1A, SCN2A, SCN8A*, *CACNA1E, CACNA1A)*, episodic ataxia (*CACNA1A*), migraine (*CACNA1A, SCN1A),* autism spectrum disorder (*SCN2A, CACNA1C*) and pain disorders (*SCN9A*, *SCN10A*, *SCN11A*). Disorders affecting cardiac (*SCN5A CACNA1C*) or skeletal muscle (*SCN4A*, *CACNA1S*), or the retina (*CACNA1F*) have also been associated with variants in these gene families (for references see Table 1). Pathogenic variants in these genes often contribute to severe early onset disorders which are less frequently passed on to the next generation. This selective pressure is captured by the depletion of functional variants in those genes in the general population (median loss-of-function observed/expected upper bound fraction of 0.29 (Karczewski et al., 2019). Beyond rare diseases and high-penetrance variants, common variants at CACNA1 or SCN loci have also been associated with highly related common disease endpoints. For example, GWAS have identified genome-wide significant SNP-associations at loci including *CACNA1C* and *CACNA1I* for schizophrenia (Ripke et al., 2014), *SCN1A* for epilepsy (ILAE et al., 2018) and *SCN10A* and *SCN5A* for atrial fibrillation (Roselli et al., 2018).

**Table 1.**
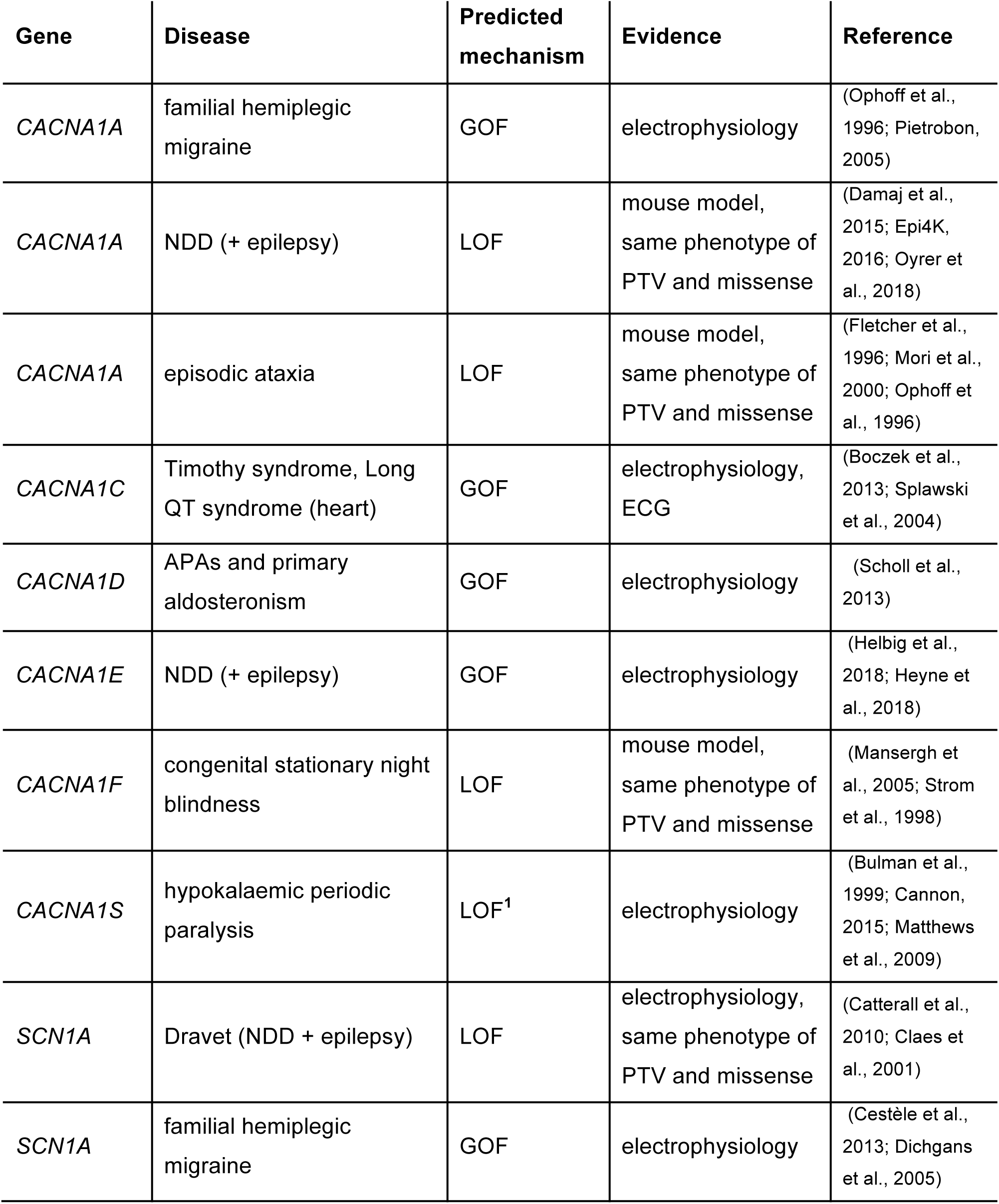
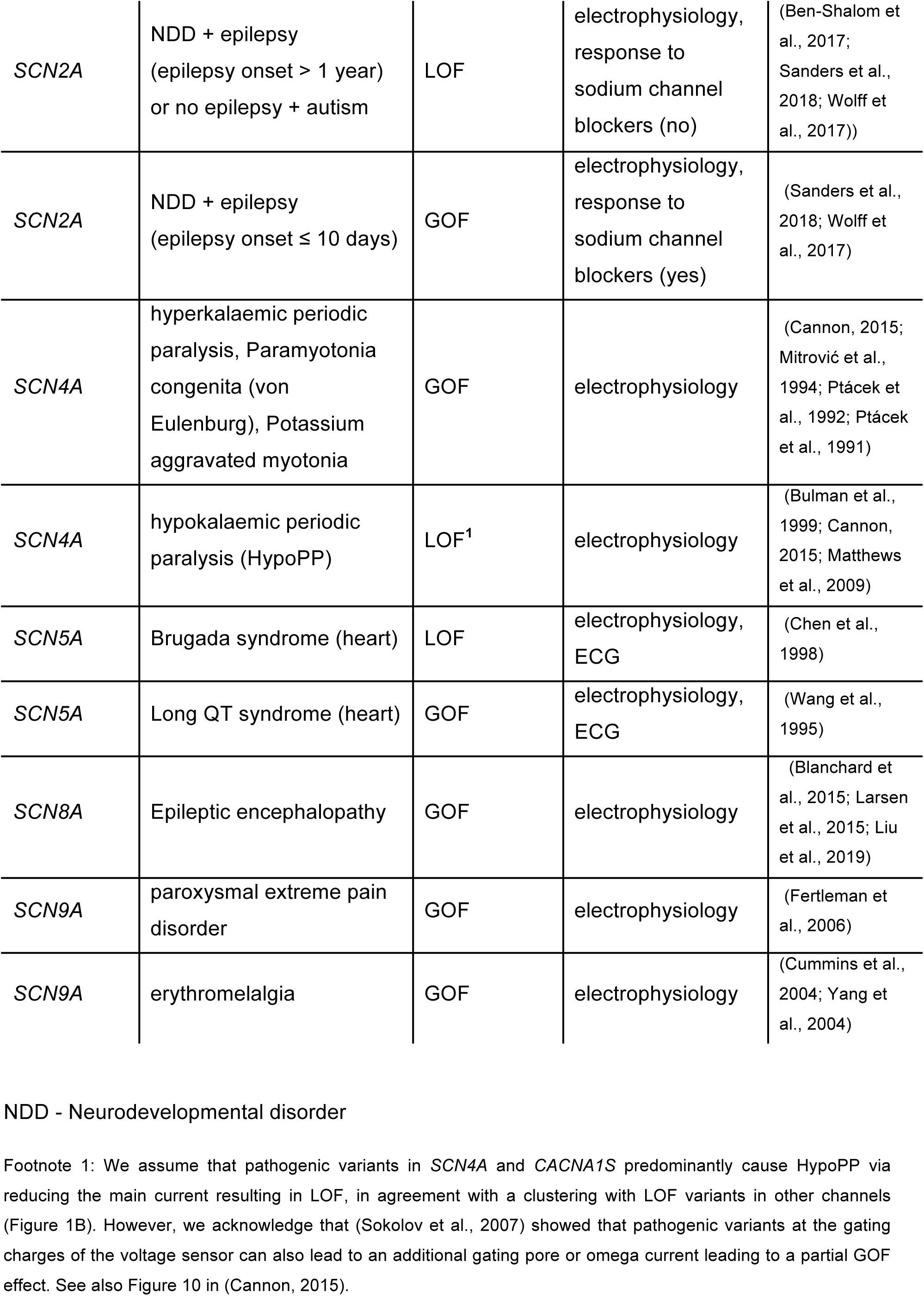
Gene – disease mechanism. This table lists references for the associations of CACNA1/SCN genes with diseases and GOF or LOF mechanisms.

Phylogenetic analyses have found that Na_v_s and Ca_v_s share a common ancestral gene (Yu and Catterall, 2004) and they have previously been defined as one gene family (Vilella et al., 2009). Na_v_s and Ca_v_s accordingly share a similar structure composed of four very similar domains I, II, III and IV, each consisting of six membrane-spanning segments S1-S6 (Catterall, 1995; Catterall and Swanson, 2015). These four domains come together, as a pseudo-heterotetramer, to form a functional channel. In the center of the structure is the pore domain which is composed of S5-S6 segments, surrounded by four voltage sensor domains (VSD) formed by S1-S4 segments. The general architecture of Na_v_s’ and Ca_v_s’ voltage-sensing and pore modules is comparable (Catterall and Swanson, 2015) and their function and structure have been extensively studied. Protein domains are associated with specific functions and diseases across channels (Brunklaus et al., 2014; Catterall and Swanson, 2015; Huang et al., 2017). It has therefore previously been suggested that mutations in similar structural domains entail similar functional outcomes in Na_v_s (Brunklaus et al., 2014), Ca_v_s (Stockner and Koschak, 2013) and Na_v_s and Ca_v_s (Moreau et al., 2014). It has also been suggested that pathogenic variants in Na_v_s and Ca_v_s occur preferentially at functionally equivalent amino acids across the gene family alignment of Na_v_s and Ca_v_s (Walsh et al., 2014). While functional differences between channels exist, particularly among calcium channels (Heyes et al., 2015), with only few amino acid (aa) changes a bacterial sodium channel was shown to be experimentally turned calcium-selective (Heinemann et al., 1992) thus illustrating the great degree of functional homology between Na_v_s and Ca_v_s. In addition, disease-associated missense variants are enriched at aa sites that are conserved across paralogs, i.e., gene family members, (Lal et al., 2017) including in sodium and calcium channels (Walsh et al., 2014; Ware et al., 2012). This further supports the hypothesis that similar biophysical pathomechanisms are involved across Na_v_s and Ca_v_s and analyzing them jointly should increase statistical power to identify disease-associated protein features. Disease-associated biological features such as protein structure and conservation metrics have been successfully used in predicting pathogenic versus neutral effects of aa changes (Adzhubei et al., 2013; Kircher et al., 2014). In voltage-gated potassium channels pathogenic variant prediction of only one gene, *KCNQ1,* (Li et al., 2017) or the K_v_ gene family (Stead et al., 2011) have been conducted with the aim of improving specificity of variant prediction in comparison to genome-wide scores. There have also been attempts to predict functional readouts using electrophysiology data in *SCN5A,* with limited success potentially due to sparse training data (Clerx et al., 2018).

Genetic variants that inactivate protein-coding genes by nonsense mediated mRNA decay such as stop-gain, essential splice, or frameshift variants have by definition loss-of-function (LOF) effects. Missense variants, however, can alter protein function in different ways. These functional alterations can be pathogenic (i.e. disease-causing), neutral (e.g. effects are small or can be compensated), or rarely beneficial. Pathogenic missense variants in SCN/CACNA1 genes can lead to disease through various changes in channel properties. These can for example affect the voltage dependence of steady-state activation or inactivation, the kinetics of the inactivation process or its recovery, ion selectivity, and other metrics that can be recorded in electrophysiological experiments (Yu and Catterall, 2004). In a simplified disease context, these variants are usually classified as having either gain- or loss-of-function (GOF or LOF) effects, depending on whether the net ion flow is increased or decreased. However, a variant may change more than one of the properties described above, with potential opposite functional effects, e.g. slowing down the inactivation process (causing GOF) on one hand and a low protein expression (causing LOF) on the other. In such cases, it may be estimated which of the functional alterations dominates to define it either as a net GOF or LOF variant, but this may also be difficult to determine. Clear LOF or GOF effects in different genes are associated with specific channelopathies. For example, in *SCN5A*, LOF variants can cause Brugada syndrome, whereas GOF variants can lead to Long QT syndrome (Kroncke et al., 2018). (All known such gene-function-disease associations are displayed in Table 1). That a pathogenic variant has a LOF or GOF effect may therefore also be inferred from disease phenotypes. In multiple genes, however, including *SCN2A* (Sanders et al., 2018) and *SCN8A* (Liu et al., 2019), phenotypic differences between LOF and GOF variants are not clear-cut or not always present at the time of diagnosis. Knowing an individual variant’s functional effect can improve prognosis, enable precision therapy (Chiron et al., 2000; Jen et al., 2007; Larsen et al., 2015; Schoonjans et al., 2015; Wolff et al., 2017) and potentially avoid incorrect treatment that could have aggravating consequences (e.g. treatment with sodium channel blockers in individuals with loss-of-function variants in *SCN2A* (Wolff et al., 2017) or *SCN1A* (Brunklaus et al., 2012)). However, current variant prediction usually only focusses on whether a variant has a disease-causing or neutral effect. We therefore introduce here a machine learning-based statistical model that can classify variants in Na_v_s and Ca_v_s as LOF, GOF or neutral thus providing a valuable resource for clinical genetics, gene discovery as well as the experimental ion channel community.

## Results

### Similar molecular mechanisms in different Na_v_s/Ca_v_s lead to LOF and GOF

Genetic variants in different Na_V_s/Ca_V_s lead to disease in diverse contexts. Comparing expression data (GTEx, 2015) and gene phenotype associations (Kohler et al., 2017), we show that tissue-specific gene expression was correlated with tissue-associated phenotypes (Figure S1) thus confirming longstanding hypotheses. For example, pathogenic variants in *SCN5A* contribute to heart diseases (Brugada Syndrome and Long QT syndrome), and the *SCN5A* encoded protein Na_v_1.5 is predominantly expressed in heart tissue. Expression in different tissues and cell types could thus explain the clinically diverse disease spectrum of Na_v_s/Ca_v_s, while allowing the possibility that similar alterations on protein structure cause heterogeneous diseases across different channels.

We therefore gathered variants in SCN/CACNA1 genes in individuals with disease from different sources (Helbig et al., 2018; Heyne et al., 2019; Heyne et al., 2018; Landrum et al., 2016; Stenson et al., 2017; Wolff et al., 2017), Table S1. We filtered these to 1521 likely pathogenic variants using ACMG-criteria (Richards et al., 2015) where possible (see Methods). Most diseases associated with Na_v_s/Ca_v_s are caused by either GOF or LOF effects. Thus, we infer whether a likely pathogenic variant has a GOF or LOF effect from disease phenotypes based on known gene-disease mechanisms. For example, in an individual with Brugada syndrome and a pathogenic variant in *SCN5A,* we assume that the variant has a LOF effect, as it has been previously described that most pathogenic *SCN5A* variants cause Brugada syndrome via a LOF mechanism (Chen et al., 1998; Kroncke et al., 2018). We screened the literature for such known gene-disease-mechanisms (for an overview, see Table 1). Applying this knowledge, from the 1521 likely pathogenic variants, we were able to classify 518 variants as likely LOF and 309 variants as likely GOF in 12 different genes across 19 diseases. 11 diseases had inferred LOF variants, 8 had inferred GOF variants. We set out to show that variants with inferred LOF or GOF effects were clustered at corresponding aa sites in Na_v_s/Ca_v_s as it would greatly boost our power to be able to jointly analyze LOF and GOF variants of different Na_v_s/Ca_v_s. In order to compare variant location of different Na_v_s and Ca_v_s, we mapped variants on a combined gene family alignment of all 20 Na_v_/Ca_v_ sequences (details see methods). We then correlated variant densities between all 19 diseases (method: Kendall correlation, Figure 1A). When variant densities of two diseases are significantly correlated, their variants are clustered at corresponding aa sites. We obtained 40 unique variant density correlations between diseases. 37 of the 40 significant correlations involved GOF variants of which 31 occurred between two diseases that were both inferred to be caused by GOF variants. This suggests that GOF variants are clustered at similar aa sites in different channels. We performed a principal component analysis to summarize all disease-disease correlations as measured by Kendall’s tau. The first principal component (PC1) perfectly separated diseases with inferred LOF from those with inferred GOF variants (Figure 1B). This indicates regional clustering of LOF and GOF variants and thus shared mechanisms lead to LOF or GOF in different ion channels. We hence combined LOF and GOF variants of Na_v_s and Ca_v_s in further analyses (see variants of all Na_v_s/Ca_v_s mapped on *SCN2A* in Figure 2).

**Figure 1.**
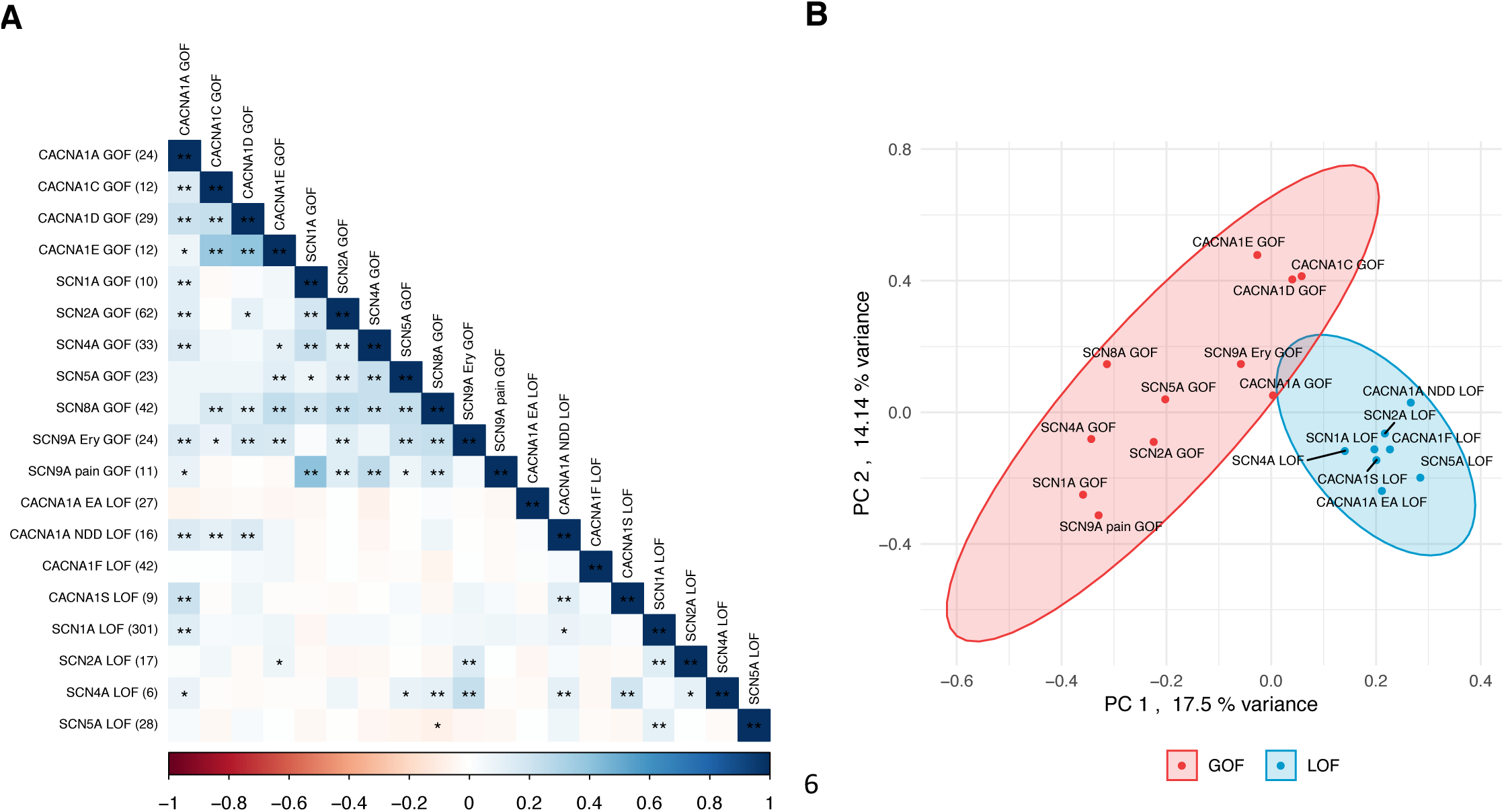
Clustering of inferred GOF or LOF variants in different genes. Inferred GOF and LOF variants were mapped on the gene family alignment of all SCN/CACNA1 genes and aa sites with gaps in the alignment removed. Variants were counted in a sliding window of 3aa. **A)** Correlation of variant densities (method: Kendall). Positive values of *tau* are blue, negatives are red (see legend). Correlations withstanding Bonferroni correction are marked with **, correlations with p-value < 0.01 are marked with *. Genes are sorted by the first principal component of the correlations (*tau*, see panel B). **B)** Principal component analysis of the correlations (*tau*). LOF variants are blue, GOF red. GOF variants in *SCN9A* are subdivided into the diseases erythromelalgia (“Ery”) and paroxysmal pain syndrome (“pain”). LOF variants in *CACNA1A* are subdivided into the diseases neurodevelopmental disorder (”NDD”) and episodic ataxia (”EA”).

**Figure 2.**
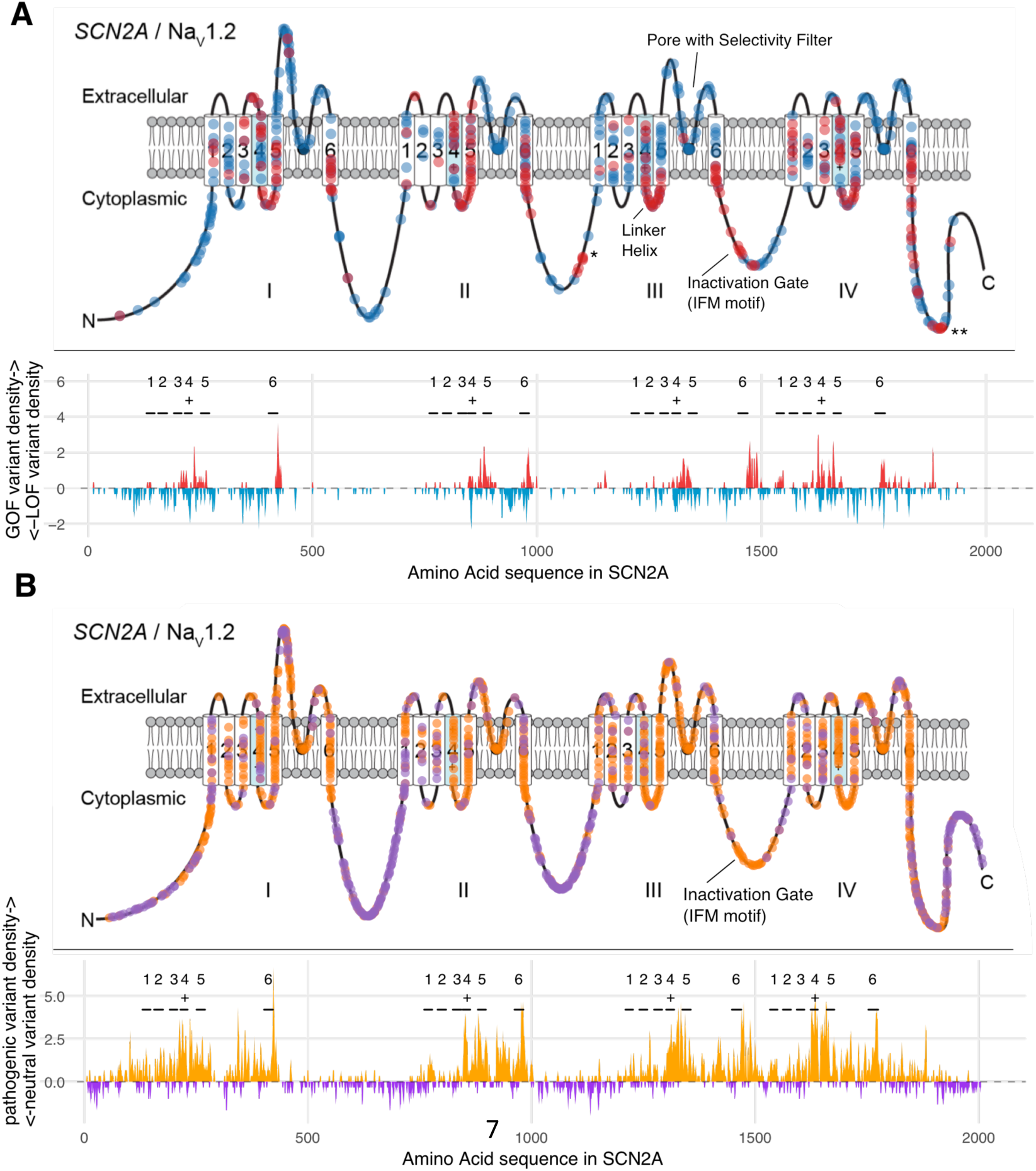
**Missense variants in SCN (Na_v_s) and CACNA1 genes (Ca_v_s) are mapped on the linear protein structure of *SCN2A*** (Sanders et al., 2018). Panel A shows inferred GOF (red) vs. LOF (blue) missense variants. Panel B shows likely pathogenic variants (orange) vs. neutral variants (purple). In both panels, upper plots show individual variants and lower plots show variant densities in a sliding window of 3aa. Na_v_s and Ca_v_s are composed of four similar domains I, II, III and IV that associate to form a channel. In each domain, transmembrane segments S1-6 are labelled with 1-6. S5-6 form the channel pore and S4 contains the voltage sensor which is labelled with “+” to illustrate the positive gating charges. The * at site 1151 refers to a cluster of GOF variants in *CACNA1C* in individuals with LongQT syndrome, ** at site 1882 refers to a cluster of GOF variants in *SCN2/8A* (see discussion). Variants with MAF > 10^-4^ in gnomAD (non-neuro) (Karczewski et al., 2019) were selected as neutral variants. Variants were inferred to be LOF or GOF from disease phenotypes (see Table 1, methods).

### Machine learning method predicts LOF vs GOF variant effects

We gathered 89 structure-based and sequence-based protein features putatively enriched for LOF versus GOF or neutral versus pathogenic effect variants. Structure-based annotations included protein secondary structure, protein’s accessible surface area and structural (e.g. cytoplasm) and functional (e.g. channel pore, selectivity filter) protein domains. Sequence-based features included conservation metrics across the 20 genes (including our own ancestry conditional selection score, see Methods), physicochemical properties of aa and deleteriousness of aa changes like “missense badness” (Samocha et al., 2017). We tested all binary protein features (features that can only take the values true or false) for an enrichment of inferred LOF, GOF, pathogenic or neutral variants with Fisher’s Exact tests (Figure 3). 6 out of 9 structure-based features and 3 out of 12 sequence-based features were enriched for 518 LOF (e.g. pore, selectivity filter) or 309 GOF variants (e.g. S4-5 linker helix, cytoplasm). In Figure 2 and Figure S2 variants are mapped on the linear sequence of *SCN2A,* in Figure S3 they are mapped onto the 3D protein structure of *SCN2A*, Figure S4 shows quantitative protein features of GOF, LOF and neutral variants.

**Figure 3.**
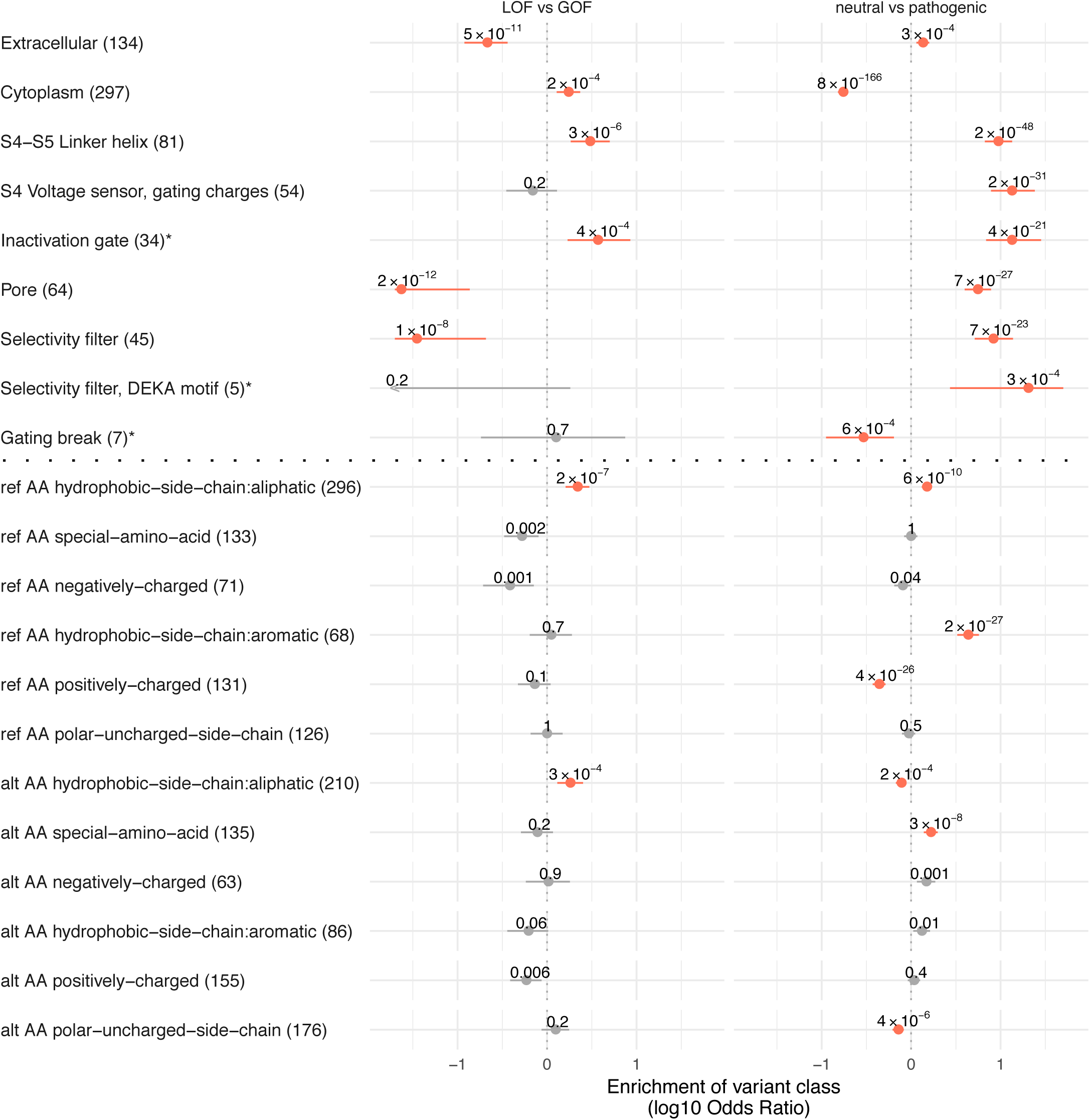
GOF, LOF and neutral variants are enriched in multiple protein features. In this figure, we show which protein features contain significantly more GOF variants than LOF variants (first column) and significantly more pathogenic than neutral variants (second column), for SCN and CACNA1 combined. Associations that are significant after Bonferroni correction for 2 x 21 tests (p-value < 0.001) are labelled red. We used Fisher’s exact tests to compare variant counts. Point estimates (log10 odds ratios) > 0 indicate a protein feature’s enrichment for GOF variants (first column) or pathogenic variants (second column). The features labelled with * are only present in Na_v_s (inactivation gate, DEKA motif of the selectivity filter) or Ca_v_s (gating break). The horizontal bars show the 95%-confidence intervals of the odds ratio point estimates that are log10-transformed and cut at -1.7 and 1.7 for clarity. Log10 odd’s ratios > 1.7 or < -1.7 are shown as arrows.

We next sought to leverage these associations of protein features with variant effects to train a prediction tool that outputs the probability that a variant results in GOF or LOF. For this, we trained a machine learning model on all 89 protein features of all 518 LOF and 309 GOF variants (Table S1). We also separately predicted neutral versus pathogenic effects (see next section). To assess the performance of our model, we set aside a test data set of 82 randomly chosen variants before the modelling process. We measured performance with the following metrics: balanced accuracy (BA), Cohen’s kappa (kappa), Matthew’s Correlation Coefficient (MCC) and Receiver Operating characteristic (ROC). The aim is usually to maximize these or similar metrics during model training. BA, kappa and MCC are performance metrics aiming to summarize a 2x2 contingency table of true positive/negative and false positive/negative predictions with a single number (Baldi et al., 2000) (see methods). The ROC curve is created by plotting the true positive rate against the false positive rate at various probabilities (Figure 4A-D). Predicting the test data of 82 variants our model reached following performance: balanced accuracy (BA) 0.80, Cohen’s kappa (kappa) 0.57, Matthew’s Correlation Coefficient (MCC) 0.59, ROC 0.85 (for ROC curves see Figure 4A-D, for performance during training see Figure S5B and C). These results indicate good predictive power when compared with other variant prediction methods (Adzhubei et al., 2013; Kircher et al., 2014; Li et al., 2017). Using this model, we ranked the relative influence of the 89 features on the prediction of LOF versus GOF effects (Figure 4E). The top two features were GOF variant density features. The ten most important features also included three different aa hydrophobicity scores, three different conservation features and the Grantham score. Our tool named “FunNCion*”* (**fun**ctional variant prediction in voltage-gated **N**a^+^/**C**a^2+^ **ion** channels) with predictions of all possible single nucleotide missense variants in Na_v_s/Ca_v_s can be found at http://funNCion.broadinstitute.org.

**Figure 4.**
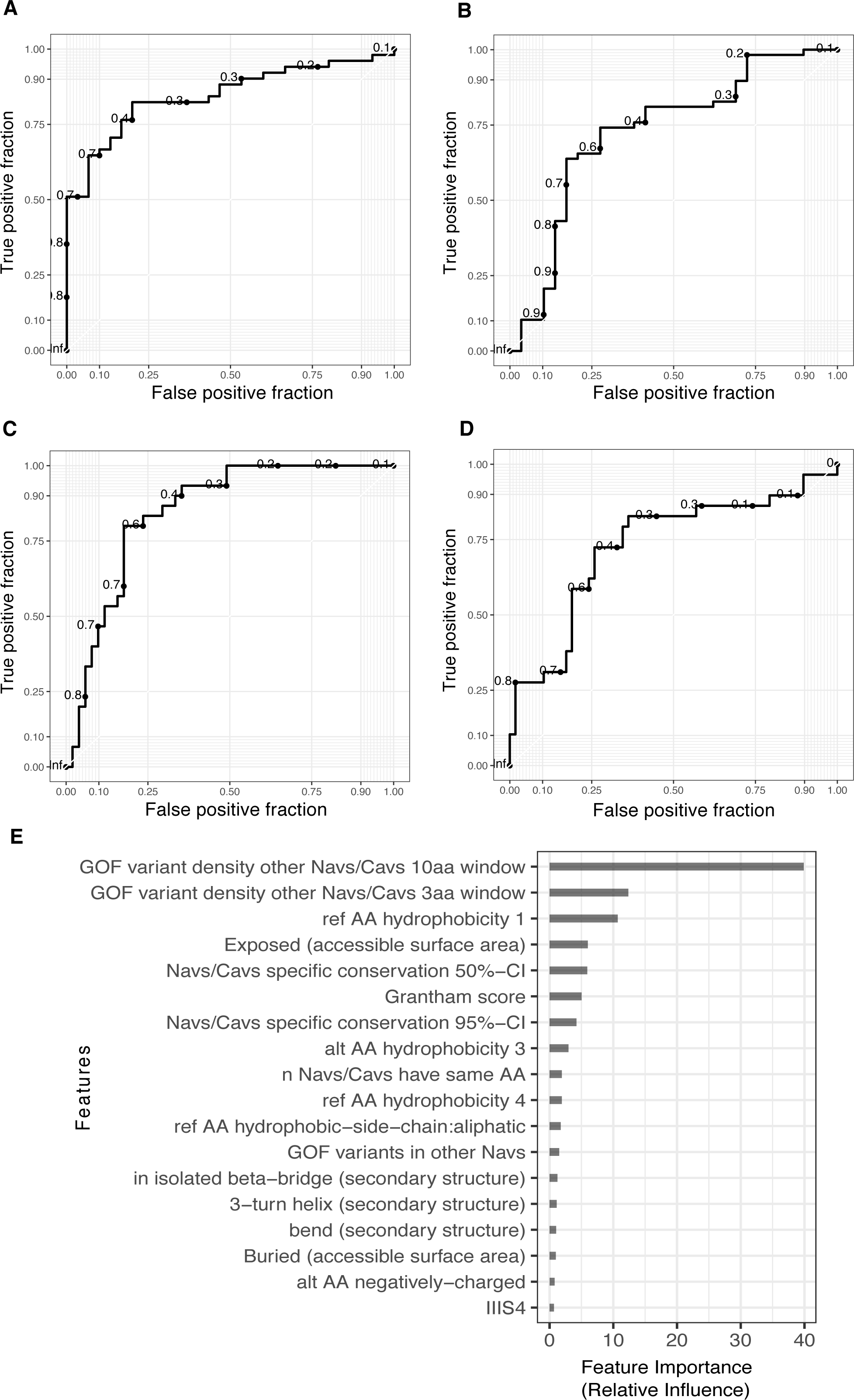
Variant prediction of LOF and GOF effects in Na_v_s and Ca_v_s. Our statistical model (method GBM) was trained on 746 variants whose functional effects were inferred from disease phenotypes. Here, we show how the model predicts LOF/GOF variant effects in two independent test data sets: 82 disease phenotypes, randomly picked from training data before model training (panels A and C) and 87 functionally tested variants, described in paragraph *Validation of funNCion with functionally tested variants* (panels B and D). **A)** prediction of LOF disease phenotypes, sensitivity=0.76, specificity=0.83. **B)** prediction of LOF electrophysiology experiments, sensitivity= 0.74, specificity= 0.72. **C)** prediction of GOF disease phenotypes, sensitivity=0.83, specificity=0.76. **D)** prediction of GOF electrophysiology experiments, sensitivity= 0.72, specificity= 0.74. The area under the Receiver Operating Characteristic curve was 0.85 for phenotype based LOF/GOF prediction and 0.73 for electrophysiology-based LOF/GOF prediction. **E)** Feature importance for prediction of GOF versus LOF. The relative influence of features on the prediction normalized to sum to 100 is computed as described in (Friedman, 2001). Of 89 features that went into the prediction, only the 18 features are shown that have a relative influence > 0.05 on the prediction.

We then asked whether modeling variants in Na_v_s + Ca_v_s jointly actually improves variant prediction over modelling Na_v_s and Ca_v_s separately. When using only the 573 Na_v_ variants during model training, prediction performance in Na_v_s was comparable to model training with all Na_v_s + Ca_v_s variants (BA 0.79, ROC 0.80, MCC 0.55 vs BA 0.80, ROC 0.85, MCC 0.58). Predicting Ca_v_s with a model only trained on 171 Ca_v_s gave however worse results compared to predicting Ca_v_s with a model trained on Na_v_s + Ca_v_s (BA 0.60, ROC 0.61, MCC 0.20 vs BA 0.79, ROC 0.88, MCC 0.58). In fact, it was better to predict Ca_v_s with a model just using Na_v_s compared to just Ca_v_s (BA 0.75, ROC 0.76, MCC 0.49). These results suggest that the increased power obtained by combining Na_v_s and Ca_v_s outweighs the differences between these channel types.

### Machine learning method predicts pathogenic vs neutral variant effects

We also set out to predict whether a variant has a ”neutral” or a potentially disease-causing (”pathogenic”) effect using the same features, the GBM method, and variants as in the functional variant prediction. We used the 1521 likely pathogenic variants described above including the 827 variants with LOF/GOF annotations plus 694 likely pathogenic variants for which we could not annotate with certainty whether they had LOF or GOF effects (see Methods). We used 2328 variants in gnomAD (Karczewski et al., 2019) in individuals who were ascertained to have no neuropsychiatric phenotypes as putative neutral effect variants. Before model training, we filtered neutral variants by frequency according to the level of genic constraint to remove rare potentially mildly deleterious variants from the neutral data set (see Methods). Similar to our functional variant prediction, we also randomly split our data set before modelling to retain 10% of variants for testing. Predicting the test data of 233 variants with our model, we obtained a BA 0.90, MCC 0.78, ROC 0.95 (for ROC curve see Figure S6A). As further validation, we predicted 89% of additional 1466 variants in genes in gnomAD that were not part of the modelling process as neutral. We predicted 466 variants in SCN genes with BA 0.86, MCC 0.64, ROC 0.93 and 1379 variants in CACNA1 genes with BA 0.93, mcc 0.32, ROC 0.97. The top 3 features with the largest relative influence on the prediction were part of a paralog-specific conservation metric “parsel” we developed for this project that estimates selection pressure while accounting for the shared evolutionary history of SCN/CACNA1 genes (see Methods). A further four conservation features were present in the top 10 features (see Figure S6B) in contrast to the LOF vs. GOF prediction dominated by variant density features (see Figure 4E). To test the performance of our model, we compared it to other popular variant pathogenicity prediction methods. To do this, we combined the two test datasets to a total of 1824 variants and removed 21 variants used in the training of PolyPhen-2. Our method performed comparably (ROC 0.95) to the three popular variant prediction tools CADD (Kircher et al., 2014) (ROC 0.79), PolyPhen-2 (Adzhubei et al., 2013) (ROC 0.85) and MPC (Samocha et al., 2017) (ROC 0.86, see Figure S6B). Pathogenicity predictions of all possible single nucleotide variants in Na_v_s/Ca_v_s can also be found at http://funNCion.broadinstitute.org.

### Validation of funNCion with functionally tested variants

To validate our model, we curated 119 functionally tested variants (96 unique variants, some tested in multiple studies) in the genes *SCN1A* (Brunklaus *et al*., in preparation), *SCN2A* (Ben-Shalom et al., 2017; Lauxmann et al., 2018) and SCN8A (Table S4) and performed functional experiments of 50 variants in CACNA1I (Table S5). In this and all subsequent validation analyses, we excluded functionally tested variants from the training data prior to modelling. In the published *SCN1/2/8A* data, out of the 119 variants 43 were GOF, 51 LOF, 13 mixed, 7 unclear, and 5 neutral. We removed 10 unique variants in individuals with benign familial infantile seizures as some variants had opposite effects in different studies (Figure S7). Our model then predicted the results of 87 electrophysiological experiments with an outcome of either LOF or GOF with a BA of 0.69, ROC 0.71 and MCC 0.38 (Figure 4, permutation p-value < 1x10^-4^). When subsetting to 57 variants that either fulfilled our phenotype/pathogenicity criteria of being included in our functional variant prediction training data or were associated with a severe phenotype, our model predicted the variants with BA 0.77, ROC 0.80 and MCC 0.54. All five variants with neutral effects were predicted to be neutral by our pathogenicity prediction method, significantly more than other functionally tested variants (Fisher’s Exact test, p-value 0.002, OR= Inf, 95%-CI 2.4-Inf). Our functional validation data also included 11 *SCN2A* variants in individuals where age of seizure onset was unavailable or outside of the cutoffs we used to infer GOF/LOF (see methods). We correctly predicted 9/11 of them emphasizing the benefit of our functional variant prediction when using phenotype as a proxy for variant function is unreliable.

We also tested our prediction on electrophysiology experiments of 50 variants in *CACNA1I* present in 12,332 individuals with and without psychiatric disease (Genovese et al., 2016) (Table S5). The functionally tested variants were present at different population frequencies. However, common variants are unlikely to have strong pathogenic effects – despite considerable efforts in GWAS, exome chip and exome sequencing, no common strong acting variants have been identified consistent with the fact that these would not be permitted by the strong selection against schizophrenia (Power et al., 2013). As our LOF- GOF prediction is trained and should hence only be used on likely pathogenic variants we sought to first predict whether variants were likely pathogenic or neutral using our own method described above. Variants that were more rare were more likely to be predicted pathogenic despite variant frequency not being a component of the model (Spearman Rank correlation between minor allele frequency (MAF) in the population cohort gnomAD (Karczewski et al., 2019) and pathogenic prediction, rho= -0.60, p-value= 3.3x10^-6^, Figure 5A). Interestingly, we found that whether a variant was predicted pathogenic correlated with whether a variant had a functional effect when considering variants that were present in only one individual (BA 0.75, ROC 0.77 and MCC 0.44). There was however no association of pathogenicity and functional effect in 19 variants that were present in >10 individuals in gnomAD (BA 0.46, ROC 0.36 and MCC -0.14). That is consistent with the above-mentioned statement, that common variants should have no strong pathogenic effects suggesting functional effects found at higher variant frequencies were likely milder or not disease-causing. Given, that a variant is pathogenic, we predict its functional effect (LOF or GOF) with BA 0.83, ROC 0.78 and MCC 0.58 (see Table S5, Figure 5B). We then combined the z-scores of four electrophysiology parameters to investigate how well we could predict variants with different magnitudes of functional effects. Firstly, pathogenicity probability positively correlated with the combined experimental z-score in variants with functional effects (Spearman correlation, p-value: 0.02, rho: 0.43, Figure 5C). In a logistic regression model, we also found that the strength of the functional effect (combined experimental z-score) influenced whether funNCion correctly predicted LOF or GOF functional effects (coefficient 0.29, p-value 0.02, Figure 5D). Accordingly, when only analyzing the ten variants with a combined z-score of four experimental parameters >=16, we predicted functional effect (LOF or GOF) with BA 0.94, ROC 0.89 and MCC 0.67. Taken together, these results suggest, we can successfully predict LOF vs. GOF in functionally tested variants, with an increased performance in variants with larger functional effects and variants that are more likely pathogenic.

**Figure 5.**
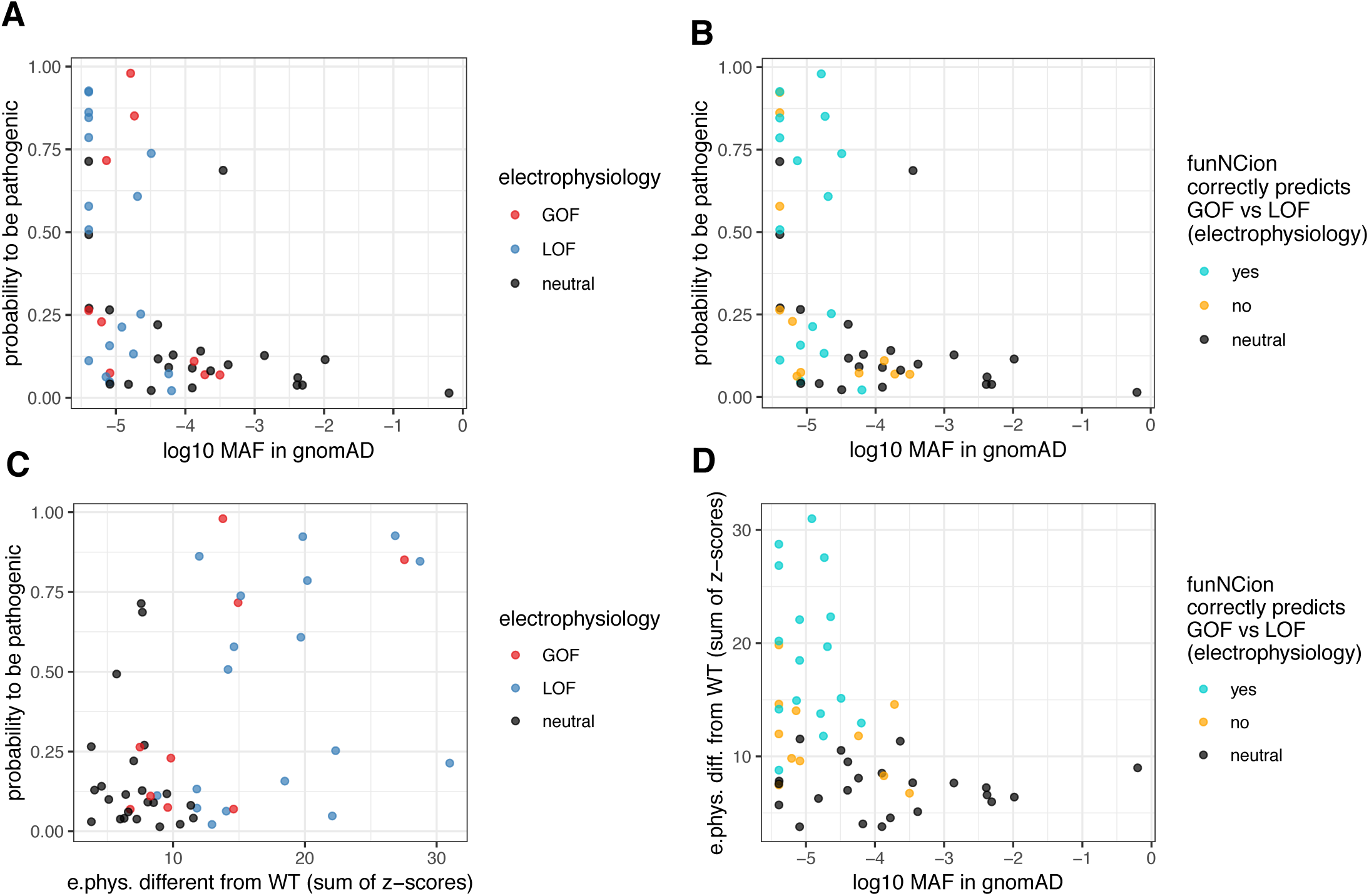
Functional and pathogenicity prediction of 50 experimentally tested variants in *CACNA1I*. This figure shows that 10 out of 12 variants that our method labelled as pathogenic also show functional effects in experiments and are rare in the general population (gnomAD). Given that a variant is pathogenic, we predict its functional effect (LOF or GOF) correctly with BA 0.83, ROC 0.78 and MCC 0.58. In panels A-C, the y-axis indicates the probability to be pathogenic. If the probability to be pathogenic is > 0.5, we label the variant as pathogenic. In panels A and D, a combined z-score indicating how much the four electrophysiology parameters differ from wildtype (see Methods) is shown on the x-axis or y-axis. In panels A, C and D, we show minor allele frequency (MAF, log10) in gnomAD on the x-axis. In panels A and B variants are labelled according to classification in electrophysiology experiment (GOF: red, LOF: blue, neutral: black). In panel C and D variants are labelled according to the agreement of functional variant prediction (LOF or GOF) with electrophysiology experiments given that they are functional (correctly predicted: turquoise, wrongly predicted: yellow, neutral variants: black).

### Validation of funNCion with large datasets of population controls and neuropsychiatric diseases

We first predicted functional and pathogenicity effects of missense variants in 114,704 individuals without severe pediatric and neurological disorders in gnomAD (Karczewski et al., 2019). We set out to test which factors predict a variant’s probability to be pathogenic (method: linear regression). The most significant predictor was -log10 MAF in gnomAD (p-value 2x10^-74^, coefficient: -0.07) i.e. pathogenic variants were at significantly lower frequencies in gnomAD. This is expected, as selection should not allow deleterious variants to rise to high population frequencies (Power et al., 2013); see *CACNA1I* in previous paragraph. We also observe this in individual genes (Bonferroni corrected p<0.0025 for 8 *CACNA1* and 5 *SCN* genes, p<0.01 for 3 genes) except *SCN7A*, *SCN10A*, *SCN11A* and *CACNA1F* (see Figure 6A). A positive predictor of variant pathogenicity was a gene’s LOEUF (loss-of-function observed/expected upper bound fraction ((Karczewski et al., 2019), p-value 2x10^-67^, coefficient: 0.21). A low LOEUF value means that the respective gene has significantly fewer protein-truncating variants (PTVs), here labelled “loss-of-function” variants as they have by definition a LOF effect, in gnomAD than expected. The equivalent value for missense variants (here termed “MOEUF”) was also significant (p-value 1x10^-4^, coefficient: 0.06). It is again expected that genes which are most intolerant to functional variants would harbor mostly neutral rather than pathogenic missense variants in a cohort of primarily healthy individuals. LOEUF being more strongly associated with pathogenicity than MOEUF suggests that Na_v_s/Ca_v_s may be generally more intolerant to loss-of-function (including LOF missense and truncating) than gain-of-function variants. To test this, we ran a linear regression model of GOF probability as a response variable. Overall, pathogenicity probability was positively associated with GOF probability (p-value 2x10^-55^, coefficient: 0.37), and LOEUF was negatively associated with GOF probability (p-value 5x10^-6^, coefficient: -0.07) while MOEUF was slightly positively associated with GOF probability (p-value 0.01, coefficient: 0.06). This is in line with the notion that most Na_v_s/Ca_v_s, but in particular those with a lower PTV tolerance, tolerate LOF missense variants less than GOF missense variants. In contrast, genes with particularly low tolerance for missense variants harbored fewer GOF than LOF variants in gnomAD (see Figure 6B). We find the association of pathogenicity and GOF probability in all individual genes (p-value < 0.0025 to correct for 20 tests) except *SCN2A, SCN8A, CACNA1A, CACNA1B, CACNA1C, CACNA1D* and *CACNA1E*. Fittingly, all of them except *CACNA1B* are implicated in severe GOF disorders and *SCN8A*, *SCN2A, CACNA1C* and *CACNA1E* have the lowest MOEUF values of all Navs/Cavs. Overall, these biologically meaningful results validate our method.

**Figure 6.**
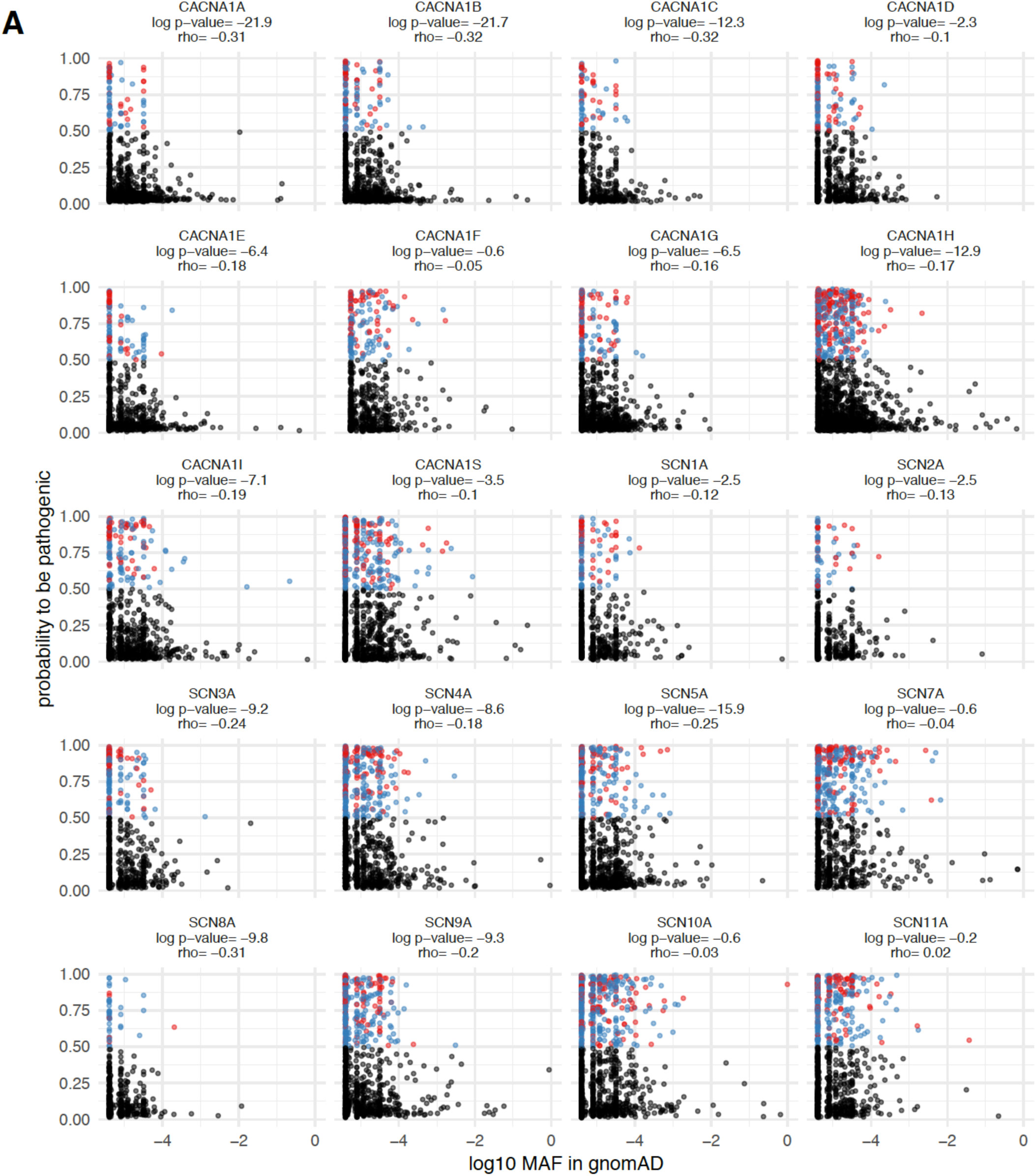
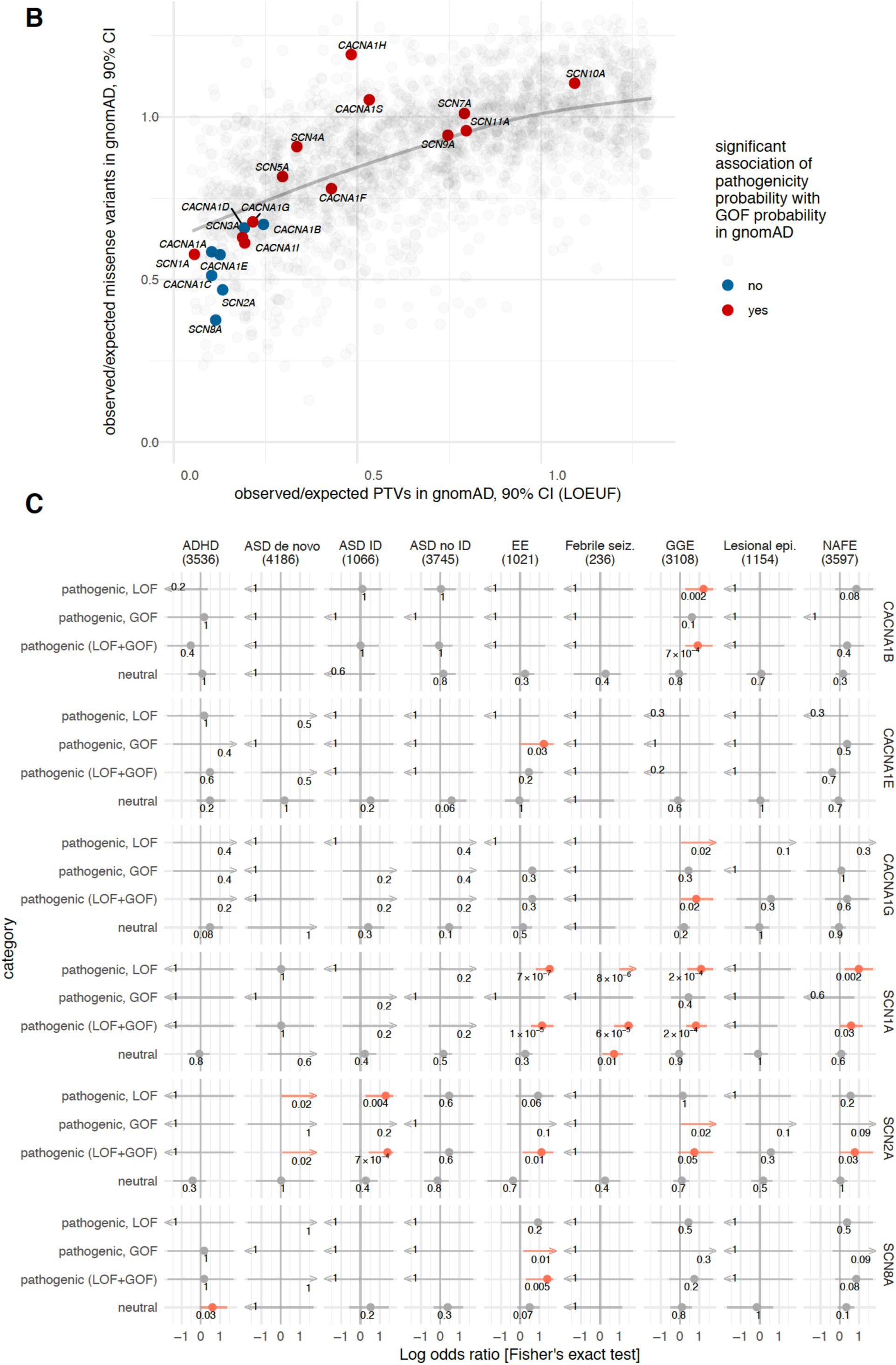
Predicting GOF, LOF, pathogenic and neutral variant effects in large cohorts of individuals with and without disease. In panel A, we show that predicted pathogenic variants (pathogenicity probability on y-axis) are at significantly lower minor allele frequencies (MAF, x-axis) in the gnomAD population cohort (rho and log10 p-value are given per gene, correlation method: Spearman, Bonferroni p-value: 0.0025). Variants predicted to be neutral are colored black. Variants predicted to be pathogenic are colored according to their predicted functional effect; GOF in blue, LOF in red. In panel B we show the 90%-CI of the observed-over-expected ratio (oe) of missense (y-axis) and truncating variants (x-axis) of SCN/CACNA1 genes in gnomAD. We plotted oe values of 3000 random genes in gnomAD in grey in the background. We colored genes red if pathogenicity probability was significantly (p-value < 0.0025 to correct for 20 tests) associated with GOF probability, potentially indicating those genes tolerate LOF missense variants less than GOF missense variants. Genes without that association (blue, p-value > 0.0025) also had the lowest oe ratios for missense variants in gnomAD (*SCN2A, SCN8A, CACNA1C,* and *CACNA1E*). In panel C we show our prediction in large datasets of individuals with diseases. We compared ultra-rare missense variants in individuals with and without epilepsy (Feng et al., 2019), ASD (Satterstrom et al., 2018a), and ADHD (Satterstrom et al., 2018a) (method: Fisher’s exact test). We found statistically significant associations withstanding Bonferroni correction (p-value threshold 8x10^-5^) only in *SCN1A.* However, nominally significant associations (orange) of known disease genes were enriched in the directions we would expect them to be (see further description in Results, Table S8). The horizontal bars show the 95%-confidence intervals of the odds ratio point estimates that are log10-transformed and cut at -1.7 and 1.7 for clarity. Odd’s ratios > 1.7 or < -1.7 are shown as arrows.

We finally tested our prediction in large datasets of individuals with and without diseases to replicate known disease associations and mechanisms. We compared numbers of ultra-rare missense variants with Fisher’s exact tests between 9170 individuals with and 8436 without epilepsy from the Epi25 Collaborative (Feng et al., 2019), *de novo* variants (DNV) in 4186 individuals with and 2179 without ASD from the autism sequencing consortium (ASC) (Satterstrom et al., 2018a); and 8347 individuals with ASD and/or ADHD to 5214 controls from the Danish bloodspot cohort (DBS)/ iPSYCH consortium (Satterstrom et al., 2018b). We found an enrichment of pathogenic LOF, but not GOF missense variants in genes, where PTVs are known to cause specific diseases. These included 29 LOF in *SCN1A* (Catterall et al., 2010) in several non-lesional epilepsies and 14 LOF in *SCN2A* in 5252 cases of autism with ID (Sanders et al., 2018) (see Figure 6C and Table S8). *CACNA1G,* a recent candidate for genetic generalized epilepsy (GGE, n=3108) was also enriched for LOF missense but not GOF variants and combining 3 LOF missense with 2 PTVs improved disease association to p=1x10^-3^. In contrast, only missense variants are known to cause EE in *SCN8A* and *CACNA1E* which were accordingly only enriched for 2 GOF missense variants in EE (p=0.01 and 0.03, respectively). We can also nominate *CACNA1B* as a potential new candidate gene for GGE. Similar to *CACNA1G*, it was enriched for 6 missense LOF variants (p=2x10^-3^) with an overall missense signal of p=7x10^-4^. Further, bi-allelic PTVs in *CACNA1B* have recently been implicated in a severe epilepsy syndrome (Gorman et al., 2019). It would therefore be plausible, that heterozygous LOF in *CACNA1B* may lead to a milder epilepsy phenotype.

## Discussion

Tailoring treatment to individual patients’ genetic variants has made significant progress in many fields of medicine in recent years (Ashley, 2016). Studying variants’ functional effects - in ion channels with electrophysiology experiments - has enabled development of precision therapies, often accelerated by repurposing existing drugs (EpiPMConsortium, 2015; Møller et al., 2019; Oyrer et al., 2018). These functional studies require considerable effort and expertise and therefore usually focus on few variants. In Na_v_s and Ca_v_s, multiple precision medicine approaches have been described (Chiron et al., 2000; Fertleman et al., 2006; Ilg et al., 2014; Møller and Johannesen, 2016; Wolff et al., 2017), however their success is dependent on the type of functional changes of pathogenic variants. Here, we present a method that predicts LOF versus GOF effects in likely pathogenic variants in SCN and CACNA1 genes - applicable across a wide range of diseases and tissues where Na_v_s and Ca_v_s are of functional relevance.

### Relation to electrophysiology experiments

In our study, we infer LOF and GOF effects of variants from disease phenotypes without functionally testing them. This poses several challenges. Phenotypes are ascertainment-biased and there is often variable expressivity of the same variant in multiple individuals. Therefore, we carefully curated our data to clearly distinguish LOF- or GOF-associated disease phenotypes.. While few variants may still be miscategorized, specifically in the large validation cohorts the *in silico* and experimental validation rates of our method suggest that enough of them were inferred correctly to successfully train a statistical model predicting LOF versus GOF probabilities with a performance similar or better than popular variant prediction tools. Further, with the goal of predicting disease-contributing variant effects, it has also some advantages to use disease phenotypes as a quasi-functional readout. While only experiments can lead to functional insight, any experimental setup constitutes a model system. Hence, a variant may have a functional effect in a lab setting, which may not always translate to a pathophysiological effect on the organismal level. A phenotype-based model considers these additional layers of complexity that *in vitro* systems are not able to reproduce. This is illustrated for example by our data for *CACNA1I*, where pathogenicity prediction correlates with functional effects only for rare variants. This is expected, as natural selection should prevent deleterious variants from rising in population frequency.

As an example, where phenotypic and functional interpretation are occasionally contradicting, we highlight the selectivity filter domain of the channel proteins. In this region, 42 out of 43 likely pathogenic variants are in individuals with LOF phenotypes including the DEKA motif in Na_v_s that conveys selectivity to Na+ ions. However, there are examples of GOF effects in electrophysiology experiments. The p.G1662S variant in *SCN10A* encoding Na_V_1.8 was implicated in small fiber neuropathy and showed GOF functionally (Han et al., 2014). However, this variant was found at a frequency of 0.0014 in the gnomAD population database including four homozygous individuals and therefore rated benign by two independent laboratories in ClinVar. Thus, the variant’s functional changes are unlikely to contribute to disease. The second one is the variant p.K1422E in *SCN2A* carried by an individual with NDD and epilepsy who was 13 months old at the onset of seizures thus corresponding to a LOF disease phenotype. In previous studies, the variant rendered the channel much elevated permeability to divalent cations like Ba^2+^ and Ca^2+^ while selectivity of Na^+^ was significantly reduced (Schlief et al., 1996). We could experimentally replicate that the variant acted as a GOF electrophysiologically in terms of permeability to Ca^2+^ (data not shown). However, we also found that the Nav1.2 p.K1422E variant carried significantly lower overall current density compared to Nav1.2 wild-type and the Nav1.2 p.T1420M isogenic variant stable cell line (Figure S8 and Methods). The current density reduction we see potentially reflects biological defects in either forward trafficking, reduced single channel conductance, increased permeability to outward Na/Cs current, or enhanced endocytosis/degradation. The apparent LOF effect in current density may override the GOF effects in Ca^2+^ influx thus explaining the overall LOF disease phenotype. These effects would be difficult to properly evaluate in transient expression systems thus illustrating the difficulty of experimentally modelling those complex proteins.

### Limitations

Our approach has several limitations. We acknowledge that the classification of variant effects into LOF and GOF oversimplifies complex electrophysiological mechanisms, even if frequently done in the literature. In *SCN9A* for example, two different types of GOF mechanisms impairing channel activation and inactivation have been shown to lead to two different diseases: erythromelalgia and paroxysmal pain syndrome, respectively (Waxman et al., 2014). With large-scale experimental electrophysiology data, it may be possible to further subdivide the GOF and LOF categories or introduce more quantitative GOF/LOF scoring systems for predictions in the future. Further, we have more functional variant and experimental validation data in Na_v_s. Therefore, predictions in Na_v_s should be more reliable than in Ca_v_s. Also, our model was trained on likely pathogenic variants in mostly severe diseases. It remains to be properly validated in individuals with milder diseases with potentially milder variant effects.

### Associations of GOF/LOF with protein’s functional regions

Our results gain important insights into which functional protein domains and motifs in Na_v_s and Ca_v_s are associated with inferred GOF or LOF effects in 827 curated likely pathogenic variants. This may provide a valuable resource for experimental follow-up studies to potentially identify new mechanistically important sites and drug targets. We can also confirm associations across diseases with much higher statistical power that have thus far been shown mechanistically or only for few pathogenic variants.

As a positive control, we recapitulate known structure-function associations such as that pathogenic variants are enriched in transmembrane segments and functionally important domains like the channel pore or inactivation machinery. As mentioned above, LOF variants were clearly associated with the ion conduction structural motifs of the pore domain (S5-S6 segments) and the selectivity filter. We confirm that the structural motifs associated with the inactivation process (Catterall and Swanson, 2015; Kellenberger et al., 1997) as well as the S4-S5 linker helix were associated with GOF variants, with the latter previously implicated in pain disorders caused by variants in *SCN9A* (Waxman et al., 2014) and in developmental and epileptic encephalopathy caused by variants in *CACNA1E* (Helbig et al., 2018) and *SCN2A* (Sanders et al., 2018). Worth noting is also a slight extension of the GOF variant cluster beyond the linker helices towards the start of S5 consistently across the four transmembrane domains. Another interesting takeaway is that GOF and LOF variants are not equally associated with transmembrane segments S1-6 at the four different transmembrane domains I-IV. This corroborates previous findings that different domains in Ca_v_s and Na_v_s have an overall structural similarity but a different contribution to the channel functioning (Chanda and Bezanilla, 2002; Pantazis et al., 2014; Savalli et al., 2016). We observe an accumulation of Na_v_ and Ca_v_ GOF variants in the cytoplasmic part downstream of each transmembrane segment S6. Exploring this further may yield interesting mechanical insights.

We highlight an accumulation of four likely pathogenic variants in *CACNA1C* encoding Cav1.2 in individuals with long QT syndrome (transcript: ENST00000347598; variants: p.P857L, p.P857R, p.R858H, p.R860G). Two of these variants were previously functionally investigated. Peak calcium currents were significantly larger in mutant channels than those of wild-type for p.R858H (Fukuyama et al., 2014) and p.P857R (Boczek et al., 2013). (Boczek et al., 2013) also identified increased surface membrane expression of the channel compared to wild type. The authors found that those variants overlapped with the so-called PEST domain (proline, glutamic acid, serine, and threonine) which is involved in protein degradation signaling, leading to increased numbers of Cav1.2 channels at the cell membrane. Interestingly, this domain as well as the cluster of GOF variants are not present in other Ca_v_s or Na_v_s pointing to a distinct GOF mechanism in Ca_v_1.2.

We also report a GOF variant cluster of nine likely pathogenic SCN variants (genes *SCN2A*, ∼*4A* and ∼*8A*). When mapped onto *SCN2A* they are located in the C-terminal domain at aa sites 1875-1887 which is in close proximity to an FGF/FHF1 binding site of a Calmodulin(CaM)-FGF complex also present in Na_v_1.4 and Na_v_1.5 (Gabelli et al., 2016). FHF1-4 interact with the C-terminal domain of Na_v_s to modulate the channels’ fast and long-term inactivation (Goldfarb, 2012). One of these variants, p.R1882Q in *SCN2A*, also showed a slower time course of inactivation (Berecki et al., 2018). Further, *de novo* GOF variants in FHF1 have been associated with epileptic encephalopathy (Al-Mehmadi et al., 2016) and variants in FHF2 with generalized epilepsy with febrile seizures plus (GEFS+). Interestingly, the C-terminal lobe of the CaM-FGF complex interacts with the conserved IQ-motif of helix α- VI of the C-terminus of all Na_v_ channels (Gabelli et al., 2016), suggesting that it may serve as an anchor for the control of activation of the channels by CaM. In contrast to the FHF binding site, the I of the potential “IQ motif” overlaps with two LOF variants in *SCN1A*. These observations could yield interesting starting points for hypotheses about this interaction.

We also identify secondary structural protein features associated with LOF and GOF variants. As expected, LOF variants are more likely to be buried in the protein where they can potentially disrupt protein stability so the probability for an aa to be buried therefore becomes a predictive feature in the machine learning model.

There exist generally more LOF than GOF variants, for *SCN2A,* a recent study estimates the incidence of LOF cases to be approximately fivefold higher than GOF cases (Sanders et al., 2018). The most important reason for this is likely that GOF can be achieved at fewer sites across the genes than LOF even though other factors like frequency of genetic testing, variant penetrance and expressivity also play a role. That GOF variants can be more easily identified by their location than LOF variants is also indicated by the fact that the two top predictors of LOF/ GOF are GOF variant density features.

### Outlook

In the future, our method could potentially be used clinically, for example to predict which individuals with pathogenic variants may be likely to benefit from a particular treatment based on their variants’ LOF or GOF effects. This may potentially be relevant in an acute clinical setting when treatment decisions must be made before functional work can be done. Examples include *SCN2A* or *SCN8A*-related epileptic encephalopathy where basically any ancillary information supporting a GOF effect would help to guide treatment. tOur method could potentially be refined with large-scale experimental data, for example introducing more types of predictions than mere LOF and GOF. Beyond individual variant interpretation, our method may also be useful to stratify drug trial cohorts into functionally meaningful subgroups of patients with variants in a given gene, especially as drug trials in rare disease are difficult to scale up. This is exemplified by two new drugs specifically targeting *SCN8A* function with a goal of rectifying the effects of a GOF variant, currently in Phase I clinical trials XEN901 (https://clinicaltrials.gov/ct2/show/NCT03467100) and GS967/PRAX-330 (https://adisinsight.springer.com/drugs/800050600). As most SCN/CACNA1 genes are depleted for functional variants in the general population it is likely that more SCN/CACNA1 genes could contribute to disease for which disease associations and/or mechanisms have not yet been elucidated. In the future, our method could therefore potentially be applied in even more diseases. Finally, our study introduces disease-phenotype-based functional variant prediction that can also be used in other genes or gene families.

## Acknowledgements

We thank the members of the Analytic and Translational Genetics Unit and Broad Institute of MIT and Harvard, specifically Kyle Satterstrom, Jack Kosmicki, Yen-Chen Anne Feng, Benjamin Neale, Tarjinder Singh, Florence Wagner and Hilary Finucane for assistance and/or helpful comments. We thank Ulrike Hedrich and Yuanyuan Liu for helpful discussion on relevant functionally characterized variants and on sharing own functional data prior to publication. We thank Johannes Krause for critical reading of the manuscript. H.O.H was supported by stipends from the German Research Foundation (DFG): HE7987/1-1 and HE7987/1-2. U.S was supported by Stiftung Charité. HL has been supported by the German Research Foundation (DFG, Research Unit FOR-2715, grants LE1030/15-1 and LE1030/16-1). We gratefully thank principal investigators and members who participated in the DBS/iPSYCH, the Epi25 consortium and the ASC consortium for making these global resources possible and available to us to validate our method. The iPSYCH project is funded by the Lundbeck Foundation (grant numbers R102-18 A9118 and R155-2014-1724) and the universities and university hospitals of Aarhus and Copenhagen. The DBS (Danish National Biobank resource at the Statens Serum Institut) was supported by the Novo Nordisk Foundation. Sequencing of iPSYCH samples was supported by grants from the Simons Foundation (SFARI 311789 to M.J.D) and the Stanley Foundation. Other support for this study was received from the NIMH (5U01MH094432-02 to M.J.D). Computational resources for handling and statistical analysis of iPSYCH data on the GenomeDK and Computerome HPC facilities were provided by, respectively, Centre for Integrative Sequencing, iSEQ, Aarhus University, Denmark and iPSYCH. The iPSYCH study was approved by the Regional Scientific Ethics Committee in Denmark and the Danish Data Protection Agency. It is made available under a CC-BY-NC-ND 4.0 International license. The Epi25 project is part of the Centers for Common Disease Genomics (CCDG) program, funded by the National Human Genome Research Institute (NHGRI) and the National Heart, Lung, and Blood Institute (NHLBI). CCDG-funded Epi25 research activities at the Broad Institute, including genomic data generation in the Broad Genomics Platform, are supported by NHGRI grant UM1 HG008895 (PIs: Eric Lander, Stacey Gabriel, Mark Daly, Sekar Kathiresan). The Genome Sequencing Program efforts were also supported by NHGRI grant 5U01HG009088-02. Additional funding sources and acknowledgment of individual patient and control cohorts are listed in Supplemental Data of the Epi25 flagship paper. The Stanley Center for Psychiatric Research at the Broad Institute supported sequencing and control sample aggregation in Epi25. The Autism Sequencing Consortium (ASC) study was supported by the National Institute of Mental Health (U01s: MH100209 (to B.D.), MH100229 (to M.J.D.), MH100233 (to J.D.B), & MH100239 (to M.W.S.); U01s: MH111658 (to B.D.), MH111660 (to M.J.D.), MH111661 (to J.D.B), & MH111662 (to S.J.S. and M.W.S.); Supplement to U01 MH100233 (MH100233-03S1 to J.D.B.); R37 MH057881 (to B.D.); R01 MH109901 (to S.J.S. M.W.S., A.J.W.); R01 MH109900 (to K.R.); and, R01 MH110928 (to S.J.S., M.W.S., A.J.W.)), National Human Genome Research Institute (HG008895), Seaver Foundation, Simons Foundation (SF402281 to S.J.S., M.W.S., B.D., K.R.), and Autism Science Foundation (to S.J.S., S.L.B., E.B.R.). Full author names and funding sources of individual cohorts contributing to the ASC can be found on the corresponding ASC flagship paper. The Genotype-Tissue Expression (GTEx) Project was supported by the Common Fund of the Office of the Director of the National Institutes of Health, and by NCI, NHGRI, NHLBI, NIDA, NIMH, and NINDS. The funders played no role in the design of the study, in the collection, analysis, and interpretation of data, or in writing the manuscript.

## Author Contributions

Conceptualization, HOH and MJD; Methodology, HOH, SI, DJP and MJD; Data Curation and Formal Analysis, HOH; Software HOH and DJP; Investigation, HOH, DBN and HRW; Writing – Original Draft, HOH; Writing –Review & Editing, HOH, DB, DJP, SI, AB, EPP, US, HL, PM, DL, AJC, JP, HRW and MJD; Resources, DBN, AB, Epi25 Collaborative, KJ, SL, JRL, RSM, EPP, US, SS, HL, PM, JP, HRW; Supervision, SI, DJP, DL, AJC, JP, HRW and MJD.

## Methods

### CONTACT FOR REAGENT AND RESOURCE SHARING

Further information and requests for resources should be directed to and will be fulfilled by the Lead Contact, Henrike O. Heyne.

### EXPERIMENTAL MODEL AND SUBJECT DETAILS

#### Automated patch-clamp

The electrophysiology experiments of 50 variants in Ca_v_3.3 and variant Na_v_1.2 p.K1422E were performed with automated patch-clamp experiments using the SyncroPatch 384PE platform (Nanion Technologies®). This automatic high-throughput patch-clamp system is able to record simultaneously up to 384 independent cells with GΩ resistance seals. The position within the chip of the different variants and wt cells was randomized to avoid artifacts due to the recording position. Cells were harvested 72h after induction for recording, as described in recent protocols for automated patch clamp for calcium channel recordings (Pan et al., 2018). Cells were rinsed with PBS (5 mL) and treated with 3 mL Accutase (STEMCELL technologies) for 5 min at 37°C, re-suspended in 10 mL of serum-free media and pelleted at 1000 rpm for 3 min at RT. The supernatant is discarded and cells are re-suspended in serum-free DMEM F12-GlutaMAX (ThermoFisher Scientific) and pECS 50% (v:v). The cells were kept until the moment of the experiment in a temperature controlled dedicated reservoir at 10° C and shaken at 200 rpm. The experiments were performed within one hour after the harvesting process. The assays were carried in single-hole chips with resistances between 4-5 MΩ after priming the chip with the following solutions (in mM), physiological extracellular solution (pECS) 10 HEPES, 140 NaCl, 5 Glucose, 4 KCl, 2 CaCl2, 1 MgCl2, 295-305 mOsm pH 7.4 (NaOH). Internal recording solution (in mM) 20 EGTA, 10 HEPES, 50 CsCl, 10 NaCl, 60 CsF, 285 mOsm pH 7.2 (by 1N CsOH). The junction potential (∼ 12 mV) and the fast capacitive component were compensated, then 15 uL of the cell suspension (50% v/v pECS/DMEM no serum) was added to each well to a final density of 50-80K cells/mL. Cell capture was promoted by holding a negative pressure of -100 mbar for 20 s. After the capture the seal was enhanced by successive hyperpolarization steps from -30 mV to -100 mV followed by the transient addition of a high Ca2+ extracellular solution (in mM) 10 HEPES, 80 NaCl, 5 Glucose, 60 NMDG, 4 KCl, 10 CaCl2, 1 MgCl2, 310 mOsm and pH 7.4 (HCl). High Ca2+ solution is washed out by successive external exchanges replacing half of the volume of the well each time with the external recording solution (in mM) 10 HEPES, 80 NaCl, 5 Glucose, 60 NMDG, 4 KCl, 6 CaCl2, 1 MgCl2, 300 mOsm and pH 7.4 (HCl). All recording solutions were prepared with ultrapure MilliQ water (18 MΩ-cm). All of the salts were purchased from Sigma-Aldrich. Hygroscopic reagents were kept in a salt desiccator container; moist salts may lead to inadequate osmolarity, a key parameter in planar electrophysiology experiments. All the solutions were filtrated with 0.22 µm PES membrane and stored at 4°C until use. The whole-cell configuration was achieved by a brief negative pressure pulse of -250 mbar. The holding potential was set at -100 mV for all the voltage protocols. Once in whole-cell configuration, the slow capacitive component (Cslow) was canceled and the series resistance (Rs) compensation was set at 80%. The data were acquired at 20 kHz and filtered at 10 kHz using Nanion proprietary software PatchControl 384 software (v.1.4.5). The data was processed on DataControl384 Version 1.5.0 previous to the analysis using the quality checkpoints along the experiment, using the seal resistance, capacitance and series resistance as qualitative parameters. The peak current and the activation steady-state parameters were obtained from currents elicited by a two-pulse protocol. During the preconditioning pulse (TP1) the cells were held for 1s to a range of voltages from -120 to 20 mV with a 10 mV increase per sweep, followed by a 200 ms test pulse at -20 mV (TP2). The peak current density value was calculated as the maximum peak current from the preconditioning pulse normalized by the capacitance. The normalized chord conductance (G/G_max_) was calculated from the I-V relationship constructed with the peak current values for each voltage during TP1, using the following equation:

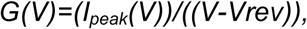

where *Vrev* is the reversal potential obtained using a linear extrapolation of the last four points of the I-V relationship mentioned above. The voltage activation process was fitted using a single Boltzmann function:

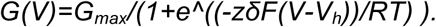

The *G_max_* was defined as the maximum value of G when the first derivative of G is minimal, *zδ* corresponds to the slope of the function and represent the voltage dependency of the activation, Vh is the voltage at which half of the *G_max_*. R, T and F refer to the gas, temperature and Faraday constant.

The steady-state inactivation parameters were obtained from the peak current values during TP2 which represent the fraction of channels able to be opened at the end of TP1. The voltage inactivation was fitted by the sum of two Boltzmann distributions according to the following equation:

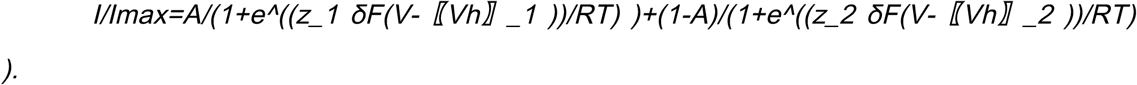

Where A represents the contribution of each individual Boltzmann to the final function normalized between 0 and 1, all the other parameters have the same meaning described above. The recovery from inactivation was obtained applying paired-pulse protocol. The protocol corresponds to a preconditioning pulse at -20 mV for 200 ms was followed by recovery inter-pulse with a variable length, ranging from 10 ms to 1600 ms. The peak current in the second pulse was normalized by the current in the first pulse, the current fraction was plotted as a function of the inter-pulse time. The data was well fitted by a mono-exponential function as follows:

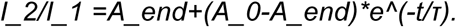

In order to compare the different variants across different parameters with different physical domains (voltage, time and current density) we transformed the different magnitudes into standard scores (Z scores), where each mean value in each parameter is represented as a standard deviation distance from the wt channel according to the following equation:

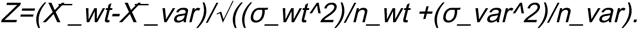

Where *X*, *σ2* and *n* stand for mean, standard deviation, and size of the sample respectively

#### Manual patch

Conventional Patch-clamp experiments under the whole-cell configuration were carried out 72h after induction. The currents traces were amplified using a MultiClamp 700B amplifier (Molecular Devices Inc.). The amplified electrical signals were acquired with a Digidata 1440A (Molecular Devices Inc.) using the pClamp10 software (Molecular Devices Inc.). Extracellular recording solution contained in mM: 2 CaCl2, 10 HEPES, 140 NaCl, pH adjusted to 7.2 with NaOH. The internal electrode was fabricated from borosilicate capilars with the puller P1000 (Sutter inst), and polished using a microforge (Narishige, Inc.) to final diameter of 1-2 um, the resistance of the internal electrode was 3-5 Mohms, when was filled with the following solution in mM: 126 CsCl, 10 EGTA, 10 HEPES, 1 EDTA, and 4 MgATP, Vjp was estimated ∼18 mV.

#### Cell lines used for investigation of Nav1.2 K1422E and T1420M mutants

Human Nav1.2 wild-type as well as Nav1.2 K1422E and Nav1.2 T1420M were introduced into FlpIn TREx 293 cell lines and selected single clone in the presence of 100 ug/mL Zeomycin and 15 ug/mL Blasticidin to generate stable cell lines. FlpIn TREx 293 cell lines were cultured and maintained in DMEM/F12 medium supplemented with 10% FBS. Cells are passaged with 0.25% trypsin at 80% confluence. The expression of Na_v_1.2 wild-type and mutants was induced by 1ug/ml Doxycycline addition to the culture for 48 hrs in T175 flasks for automated patch-clamp recording.

#### Interpretation of variants’ functional changes as LOF and GOF effects

Functionally tested *CACNA1I* variants were defined as LOF or GOF according to how the different parameters influence the Ca2+ influx during simulated afterhyperpolarization (AHP) rebound in (Andrade et al., 2016). Only variant effects with Z scores significantly higher than observed in wildtype negative controls were interpreted as functional changes. Leftward shifts in the activation midpoint, rightward shift in the inactivation midpoint, increases in the peak current density were interpreted as GOF. Rightward shifts in the activation midpoint, leftward shift in the inactivation midpoint and decreases in the peak current density were defined as LOF. When two or more parameters were affected, the preponderant functional outcome was the effect in the Ca2+ influx of the different parameters combined during the AHP rebound. A combination of opposite effect parameters was interpreted as “neutral” predictions.

#### METHOD DETAILS

#### Variants used in variant prediction

#### Likely pathogenic variants

Assuming that diseases caused by variants in SCN/CACNA1 are caused by an ultimate GOF or LOF of Na_v_s/Ca_v_s, we inferred LOF or GOF effects of likely pathogenic variants from disease phenotypes from in total 19 different diseases (see Results section for examples and Table S1 for all disease mechanisms with references). We curated published variants from ClinVar (Landrum et al., 2016)(version 05/2018), Human Gene Mutation Database ((Stenson et al., 2017) HGMD, version 05/2018), (Heyne et al., 2019; Heyne et al., 2018), and curated variants in genes *CACNA1D, CACNA1A* and *SCN2A* from the literature including unpublished phenotype data for *SCN2A* resulting in 303 inferred GOF and 524 inferred LOF variants. We only included variants for which we could infer LOF or GOF effects with high confidence. In individuals with pathogenic variants in *SCN2A*, pathogenic variants have previously been suggested to have a LOF effect for individuals with seizure onset > 3 months and GOF effect for seizure onset < 3 months (Ben-Shalom et al., 2017; Wolff et al., 2017). However, exceptions to those cutoffs have been reported (Lauxmann et al., 2018). In *SCN2A,* we therefore classified variants in individuals with seizure onset ≤ 10 days as GOF due to a clear accumulation of 93 individuals with seizure onset in the first 10 days of life. We classified variants in individuals with seizure onset > 1 year as LOF. We used criteria based on ACMG guidelines (Richards et al., 2015) to classify variants as likely pathogenic wherever possible. For example, if a variant is *de novo* and not present in controls it can be classified as “likely pathogenic” according to ACMG criteria PS2 and PM2. To increase specificity for variants in HGMD and ClinVar, we only used high-confidence likely pathogenic variants (in ClinVar corresponding to Review Status “criteria provided” or “reviewed by expert panel”, in HGMD corresponding to “high confidence”). We only included variants annotated as “Likely pathogenic” or “Pathogenic” in ClinVar and “Damaging Mutation” in HGMD. We curated HGMD and ClinVar phenotypes while also providing the original phenotype annotation for reproducibility (see Table S1). We applied a MAF filter to all disease variants. Disease-specific MAF thresholds were inferred from disease prevalence, inheritance mode and penetrance (see Table S2) as described in (Whiffin et al., 2017) using the authors app (www.cardiodb.org/allelefrequencyapp). In *SCN5A*, we only used pathogenic variants with penetrance of at least 0.7 as described in (Kroncke et al., 2018). For training of pathogenic versus neutral variant prediction we used the same variants as in the functional variant prediction in addition to 694 likely pathogenic variants for which we could not infer LOF or GOF mechanisms.

#### Neutral Variants

We used 3794 variants in the Genome Aggregation Database (gnomAD) (Karczewski et al., 2019) as variants with inferred neutral effects (see Table S2) during training of pathogenic versus neutral variant prediction. We excluded neuropsychiatric disease cohorts from the gnomAD data. We only included variants in genes where pathogenic as well as neutral variants were available in order to upsample to the same number of pathogenic and neutral variants per gene before training. We filtered neutral variants for MAF according to genic constraint with the rationale that mildly constrained genes might tolerate pathogenic noise-introducing variants at higher frequencies than very constrained genes. Specifically, we excluded singletons in all genes and excluded variants with MAF < 10^-4^ for mildly constrained genes as indicated by pRec > 0.9 and pLI < 0.9 and missense-z < 3.09. We also excluded variants with MAF < 10^-4^ in *SCN5A*, as the associated cardiological diseases have a high prevalence of up to 1 in 2000 individuals (see Table S3), and variants often have reduced penetrance (Kroncke et al., 2018).

All variants were annotated with Variant Effect Predictor (VEP) (McLaren et al., 2016) of Ensembl GRCh37 release 94. For comparisons with other variant prediction methods we used CADD, version v1.4 (Kircher et al., 2014), PolyPhen-2, version 2.2.2 (Adzhubei et al., 2013) and MPC (Samocha et al., 2017). Prior to these calculations variants used for training in Polyphen-2 were removed.

#### Features used in variant prediction

#### Variant density in other paralog genes

We performed gene family alignments of the UniProt (UniProt Consortium, 2018) canonical isoform sequence with the program MUSCLE (Edgar, 2004). We then mapped all variants in a functional category of either LOF, GOF or neutral onto the gene family alignment (family alignment files deposited at GitHub). For use as feature in our prediction, we mapped variants in functional categories back to the individual genes. We then counted variants in a sliding window of 3 and 10 aa, respectively, to account for different-sized windows of local sequence context. We used different ways to avoid overfitting in the pathogenicity and functional LOF vs GOF prediction. In the pathogenicity prediction, we computed variant densities for each gene only with variants in other genes, as we could sample to the same number of variants per gene and functional category before training. In the LOF vs GOF prediction, we could not sample to the same number of variants per functional category and gene in the training data as most genes did not have GOF as well as LOF variants. Therefore, we calculated GOF and LOF variant density on half of the training variants and used those variant densities as covariates during model training with the other half of the data.

#### Protein-based features

The experimentally solved 3D protein structures for Na_v_s/Ca_v_s are only available in parts or not of all genes (Pan et al., 2019). We therefore computationally predicted the 3D structures of all 20 channels using experimentally solved homologous protein structures from the Protein Data Bank (PDB, http://www.wwpdb.org) with the raptorX web server (Källberg et al., 2012) (see GitHub depository for p-values and scores of protein model predictions). Using the same web server, we collected the predicted likelihood of an aa residue being buried, medium and exposed (accessible surface area), and the likelihood of a residue conforming one out of the eight secondary structural types for all Na_v_s/Ca_v_s. We extracted protein structural features such as transmembrane, cytoplasm etc. from UniProt (UniProt Consortium, 2018) version 10/2017. We inferred the functionally important motifs in 3D protein structures linker helix (AA sites between transmembrane segments S4 and S5 (Catterall and Swanson, 2015)) and voltage sensor (positively charged aa K and R in transmembrane segment S4 (Catterall and Swanson, 2015)) for all proteins from the literature. We annotated the inactivation gate from *SCN2A* (aa sites 1472 to 1522) and the gating break from *CACNA1I* (aa sites 401 to 460) and mapped them via the gene family alignment to all Na_v_s or Ca_v_s, respectively. We annotated the selectivity filter from *SCN2A* (aa sites 378 to 389, 936 to 947, 1416 to 1427 and 1708 to 1719) and mapped it via the gene family alignment to all Na_v_s and Ca_v_s.

#### Amino acid-based features

We annotated physicochemical features of reference and alternative aa’ side chains by classifying them into the different physiochemical groups annotated in www.sigmaaldrich.com/life-science/metabolomics/learning-center/amino-acid-reference-chart.html. We considered Cysteine (with a reactive sulfhydryl group) and the “unique” aas (Proline and Glycine) together as “special aas”. Additionally, we used measures of aa deleteriousness such as the “missense badness” annotation from (Samocha et al., 2017), which quantifies tolerance of different types of missense aa exchanges in the general population by comparing expected versus observed numbers of all possible aa exchanges in ExAC (Lek et al., 2016). We included also the aa deleteriousness metrics “Grantham score” (Grantham, 1974) and BLOSUM score (Henikoff and Henikoff, 1992). All features of all potential aa changes have been deposited at github.com/heyhen/funNCion.

#### Ancestry conditional site-specific selection score

We constructed several features that estimate conservation within and between Na_v_s and Ca_v_s that were particularly informative in pathogenic versus neutral variant prediction. First, we simply counted how many genes have the same reference aa as the consensus sequence in the gene family alignment, similar to the parazscore (Lal et al., 2017). Secondly, we constructed estimates of ancestry informed selection pressure to explicitly account for the shared evolutionary history of paralog genes. We first estimate the ancestry of paralogs using the gene family transcript alignment using RAxML (Stamatakis, 2014) with the GTRGAMMA model. We use the best maximum likelihood tree across 100 bootstrap searches to define a strict prior on the ancestral history of paralog sequence alignments. Conditional on this tree structure and alignment, we estimate parameters of a codon model of sequence evolution (Goldman and Yang, 1994) modified to incorporate indels (Wilson and McVean, 2006) using MCMC. MCMC moves are as described previously (Palmer et al., 2017; Wilson and McVean, 2006), with ancestry conditional likelihoods evaluated using Felsenstein’s tree-pruning algorithm (Felsenstein, 1981). Code implemented in C, which we call “parsel” (**par**alog **sel**ection) and all sequences used for development of the score are freely available at github.com/astheeggeggs/parsel under the GNU General Public License, v2.

#### Differences between canonical transcripts and canonical isoforms

One Table deposited at github.com/heyhen/funNCion includes genomic coordinates of all possible single nucleotide changes that can lead to all possible missense variants in SCN/CACNA1. To maintain compatibility with UniProt protein features we only included those transcripts whose aa sequence was 100% identical to the respective UniProt canonical isoform. In 14 out of the 20 genes, this was the Ensembl canonical transcript. The two genes for which no Ensembl or RefSeq transcript mapped to the canonical isoform were *CACNA1A* and *CACNA1C*. To maintain compatibility with the genomic coordinates, we decided to use the canonical transcripts as reference, which were the best-matching transcripts for both genes. We introduced a single aa deletion of G at aa 419 and a two-aa QQ insertion after aa 2312 in *CACNA1A’s* uniprot canonical isoform to align UniProt-derived features with the canonical transcript. In *CACNA1C*, the first 1863 aa are identical in the UniProt and Ensembl canonical transcripts, where all UniProt transmembrane features are located.

#### QUANTIFICATION AND STATISTICAL ANALYSIS

Statistical analyses were done with the R and the C programming languages. We used the R package caret for most machine learning-related functions and packages ggplot and plotROC for plotting.

#### Comparison of tissue - associated phenotypes and tissue expression

We calculated similarity of genes’ HPO terms (Kohler et al., 2017) available at https://hpo.jax.org, July 2018 release) to HPO terms that are specifically associated with a given tissue using Resnik’s method as implemented in the R package ontologySimilarity. We extracted daughter HPO terms using the R package ontologyIndex. Per tissue, we correlated genes’ tissue expression with genes’ similarity with tissue-associated phenotypes using Spearman rank correlation. P-values of correlations within different tissues were combined with Fisher’s method.

#### Clustering of inferred LOF and GOF variants of Na_v_s and Ca_v_s

In order to compare variant location between all Na_v_s and Ca_v_s, we mapped the aa sites on a combined gene family alignment of all 20 Na_v_/Ca_v_ sequences (details see method’s paragraph “variant density in other paralog genes”). We then removed alignment gaps obtaining 1268 aa sites mappable to all sodium and calcium channels (61% of their canonical isoforms length of 2064 aa ± 222 [mean ± SD]). 726 of all 827 LOF/GOF variants could be mapped on to the 1268 family-aligned aa sites. We then counted the LOF or GOF variant density on the 1268 family-aligned sites in sliding windows of 3 aa hereby considering LOF or GOF effects of variants’ directly neighboring aa sites.

#### Machine learning based prediction of GOF vs LOF and pathogenic vs neutral variant effects

We used a table of all 89 protein features by all 827 variants (Table S1) to train a prediction tool that outputs the probability that a variant results in GOF or LOF. We used the R package caret’s train function to evaluate, using a 10-fold cross-validation resampling, the effect of model tuning parameters on performance and to choose the optimal parameters for the final model. Prior to resampling, we up-sampled the data to the same number of LOF versus GOF or pathogenic versus neutral variants, respectively and also to the same number of variants in SCN or CACNA1 genes. Before training of the pathogenicity model, we additionally up-sampled to the same number of variants per gene. Prior to model training, we randomly split our dataset to retain 10% of variants as a test data set for validation. We used all available features relying on inbuilt feature selection algorithms. We included whether a gene was a Ca_v_ or Na_v_ as covariate in our model, however we did not include the individual gene as covariate. We computed GOF and LOF variant density slightly differently for LOF vs GOF and pathogenic vs neutral prediction (see method section “*Variant density in other paralog genes”*). Comparing different machine learning methods, the decision-tree based algorithms Random Forest and Stochastic Gradient Boosting, also known as Gradient Boosting Machine (GBM) outperformed logistic regression, eXtreme Gradient Boosting and support vector machine (t-tests, Bonferroni-corrected p-values 2x10^-5^ to <2x10^-16^, see Figure S5A). GBM performed best so we chose GBM as the final method for our prediction model with standard parameters. During the modelling process, tuning parameter ’shrinkage’ (how quickly the algorithm adapts) was held constant at a value of 0.1. Parameter ’n.minobsinnode’ (minimum number of training set samples in a node to commence splitting) was held constant at a value of 10. Maximizing accuracy was used to select parameters for the optimal model. The final values were: number of tree iterations = 50, interaction.depth (complexity of the tree) = 1, see also Figure S5.

#### Performance metrics of machine learning based prediction

We used following measures to assess the performance of our model: balanced accuracy (BA), Matthew’s Correlation Coefficient (MCC), Cohen’s kappa (kappa) and the Receiver Operating Characteristic (ROC). Using true positives (TP), true negatives (TN), false positives (FP), and false negatives (FN) of the 2x2 contingency table of LOF vs. GOF or pathogenic vs. neutral predictions, those metrics are defined as follows (Baldi et al., 2000).

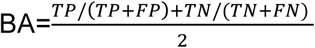

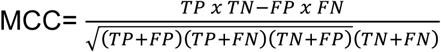

The Kappa Statistic compares the accuracy of the prediction to the accuracy of a random prediction.

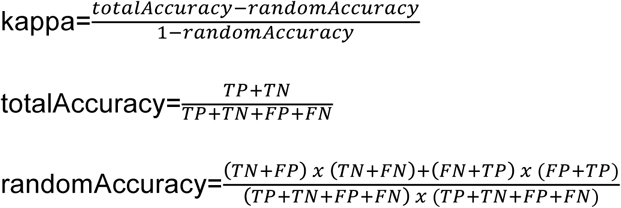

ROC is the area under the curve which is created by plotting the true positive rate against the false positive rate at various probabilities.

#### DATA AND SOFTWARE AVAILABILITY

#### Supplementary Tables

**Table S1:** All likely pathogenic variants used in functional and pathogenic variant prediction.

**Table S2:** All inferred neutral variants used in pathogenic variant prediction.

**Table S3:** Minor allele frequencies cutoffs of different diseases.

**Table S4:** Functional and pathogenic variant prediction of functionally tested variants in *SCN1A/ SCN2A/ SCN8A*

**Table S5:** Functional and pathogenic variant prediction of functionally tested variants in *CACNA1I*

**Table S6:** Prediction of variants in diseases (autism, ADHD, epilepsy)

PDB files to load 3D model of variants mapped on SCN2A (Figure S3) and homology models used to compute protein features into PyMOL: https://github.com/heyhen/funNCion

Functional (LOF vs GOF) and pathogenic variant prediction of all possible single nucleotide genomic changes that can lead to aa changes in Na_v_s and Ca_v_s:

https://github.com/heyhen/funNCion

http://funNCion.broadinstitute.org

The R code used to perform functional variant prediction and associated data tables: https://github.com/heyhen/funNCion

The C code used to perform ancestry conditional site-specific selection and associated data tables: https://github.com/astheeggeggs/parsel

ClinVar public repository (Landrum et al., 2016) provides further information for ClinVar variant identifiers referenced in Table S1: https://www.ncbi.nlm.nih.gov/clinvar/

#### ADDITIONAL RESOURCES

Please find functional and pathogenic predictions of all possible variants in Na_v_s and Ca_v_s, available as genomic or protein coordinates, on our website http://funNCion.broadinstitute.org.

