## Supplemental Information for "Predicting Functional Effects of Missense Variants in Voltage-Gated Sodium and Calcium Channels"

#### Figure S1

**Expression vs phenotypes.** It has previously been suggested, that similar mechanisms cause disease across different $Na_v$s and $Ca_v$s (Walsh et al., 2014). However, SCN/CACNA1 - associated diseases involve many different organ systems such as heart, brain, muscle etc. We set out to confirm, that these different phenotypes are largely caused by differences in gene expression and compared comprehensive expression data in 54 tissues (GTEx Consortium, 2015) of SCN/CACNA1 genes with the phenotype terms they are associated with as annotated in the human phenotype ontology. **A)** Heat map of gene expression of 10 SCN and 10 CACNA1 genes in 53 tissues (data: GTEx Consortium, v6). Expression is shown as normalized log2+1 RPKM per tissue and gene, so positive values mean higher expression than the mean and negative values mean lower expression than the mean. The figure was generated with the webtool FUMA, v1.2.8. **B)** In this panel, we calculated similarity of genes' HPO terms (Kohler et al., 2017) to HPO terms that are associated with a given tissue (Resnik's method, see Methods for details). **C)** We then correlated expression in six different tissues from panel A with HPO similarity from panel B. Across tissues, gene expression was correlated with tissue-associated phenotypes (combined p-value with Fisher's method: 0.001).

**A)**

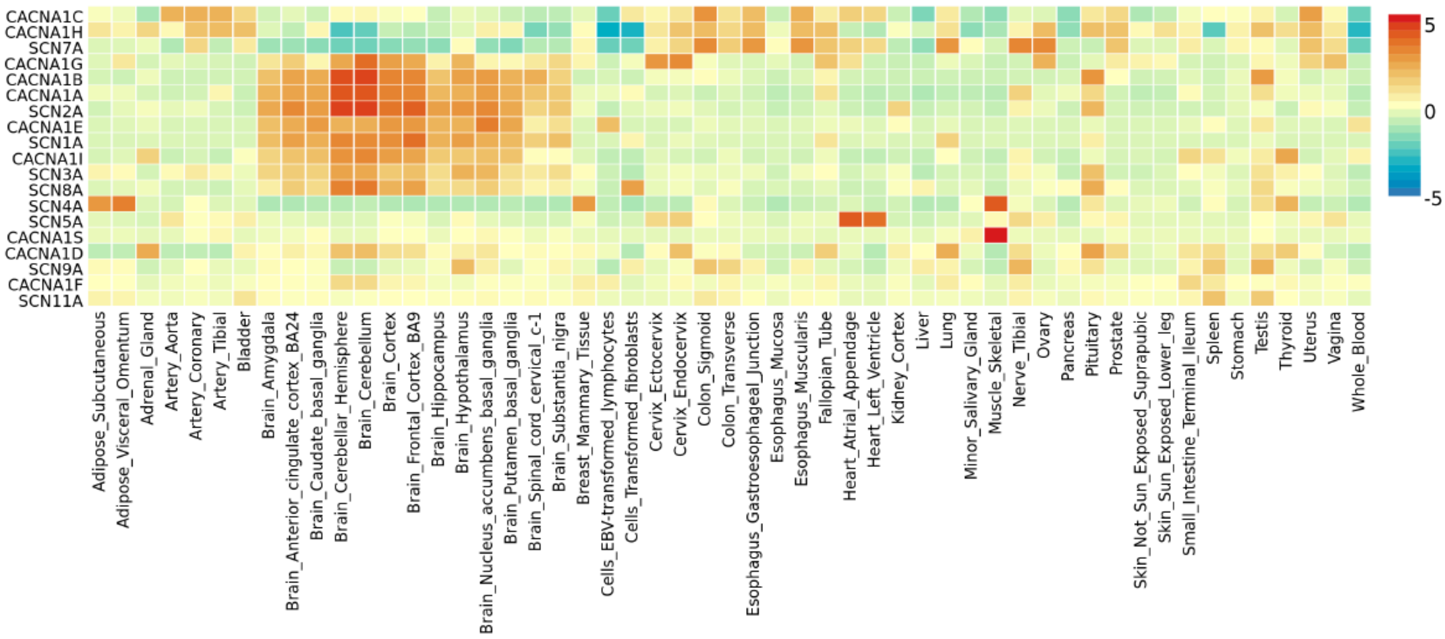

B)

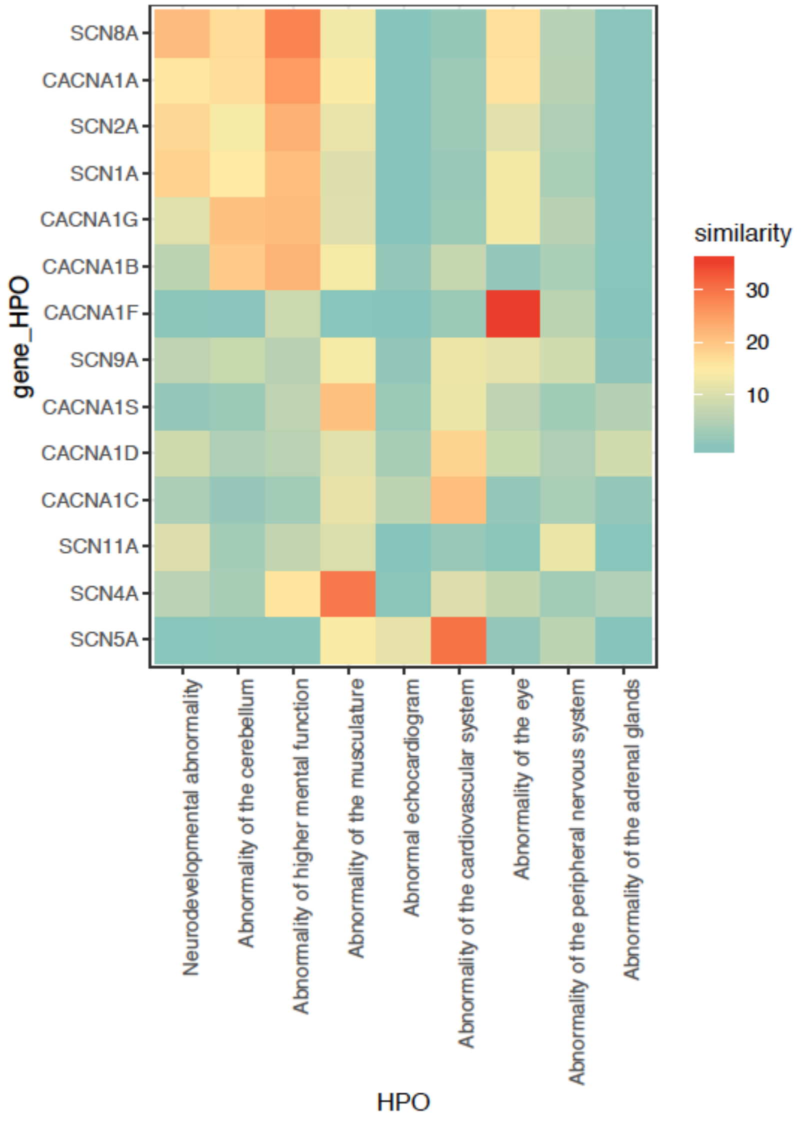
SCN8A
CACNA1A
SCN2A
SCN1A
CACNA1G
CACNA1B
CACNA1F
SCN9A
CACNA1S
CACNA1D
CACNA1C
SCN11A
SCN4A
SCN5A
gene_HPO
Neurodevelopmental abnormality
Abnormality of the cerebellum
Abnormality of higher mental function
Abnormality of the musculature
Abnormal echocardiogram
Abnormality of the cardiovascular system
Abnormality of the eye
Abnormality of the peripheral nervous system
Abnormality of the adrenal glands
HPO
similarity
30
20
10

C)

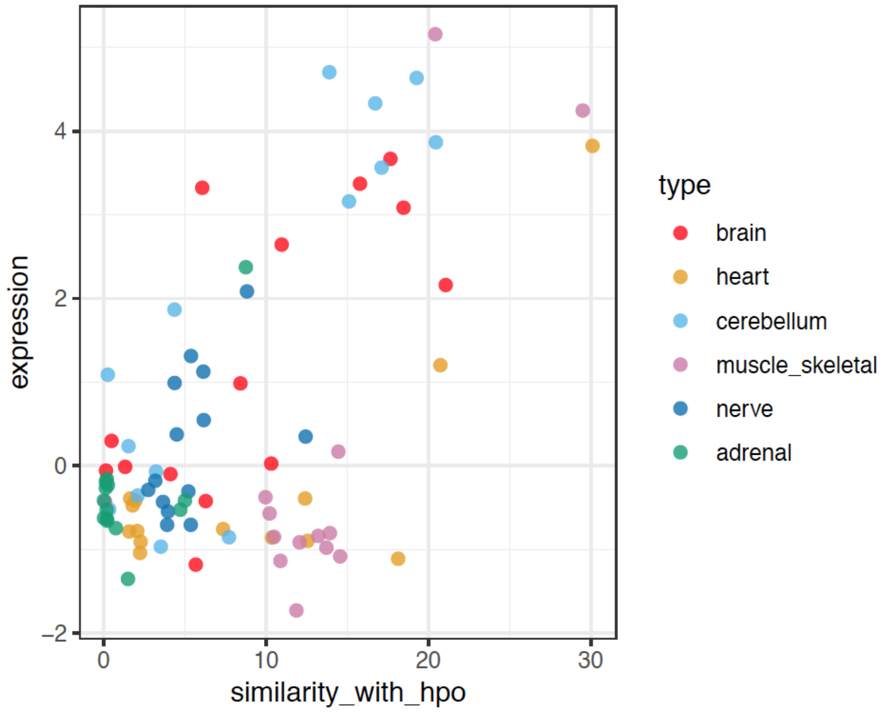
expression
4
2
0
−2
0
10
20
30
similarity_with_hpo
type
brain
heart
cerebellum
muscle_skeletal
nerve
adrenal

### Figure S2, Related to Figure 2

**Inferred GOF and LOF missense variants in SCN genes ($Na_V$s) and CACNA1 genes ($Ca_V$s) are mapped on the linear protein structure of *SCN2A.*** The upper plot shows individual variants, the lower plots show variant densities in a sliding window of 3aa. To highlight sites enriched for GOF or LOF variants, LOF variant density was subtracted from GOF variant density. GOF in both SCN and CACNA1 are shown in red, GOF in SCN in orange, GOF in CACNA1 in pink. LOF in both SCN and CACNA1 are shown in darkblue, LOF in SCN in cyan, LOF in CACNA1 in green. $Na_V$s and $Ca_V$s are composed of four similar domains I, II, III and IV, that associate to form a channel. In each domain, transmembrane segments S1-6 are labelled with 1-6. S5-6 form the channel pore and S4 contains the voltage sensor which is labelled with "+"to illustrate the positive gating charges. The * at site 1151 refers to a cluster of GOF variants in *CACNA1C* in individuals with LongQT syndrome, ** at site 1882 refers to a cluster of GOF variants in *SCN2/8A* (see discussion). Variants were inferred to be LOF or GOF from disease phenotypes (see Table 1, methods). The protein structure was taken from (Sanders et al., 2018).

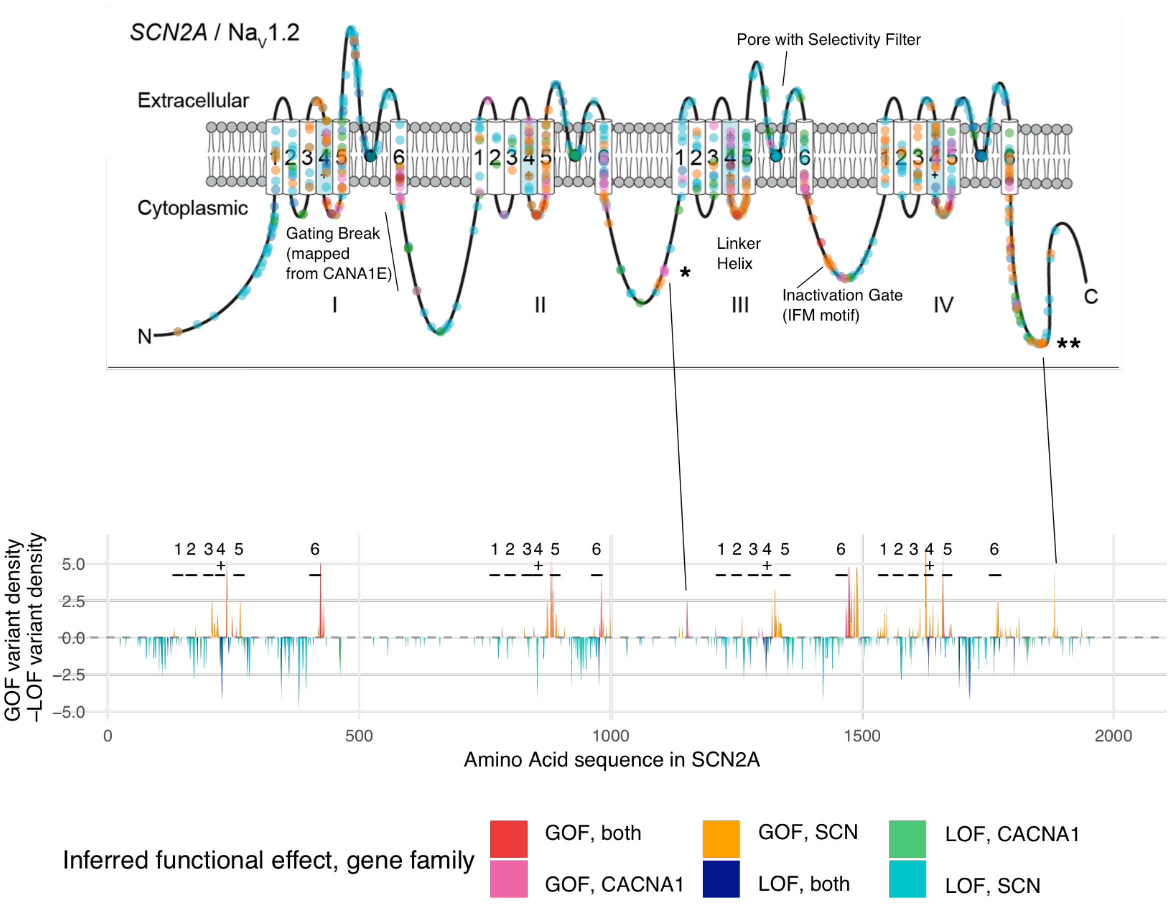

#### Figure S3, Related to Figure 2

**Pathogenic, neutral, GOF and LOF variants in SCN and CACNA1 genes mapped on the protein structure of *SCN2A*.** Panel A shows neutral variants (purple), B shows pathogenic variants (brown), C shows GOF (red) and D LOF (blue) variants. The variants were mapped on the recently published 3D structure of *SCN2A* (Pan et al., 2019)(PDB ID: 6j8e). To view the structure from different angles, we also provided a file that can be opened with PyMOL on https://github.com/heyhen/funNCion. In the PyMOL file we also provide two *SCN2B* chains in the structure (chain C in wheat and chain D in light pink).

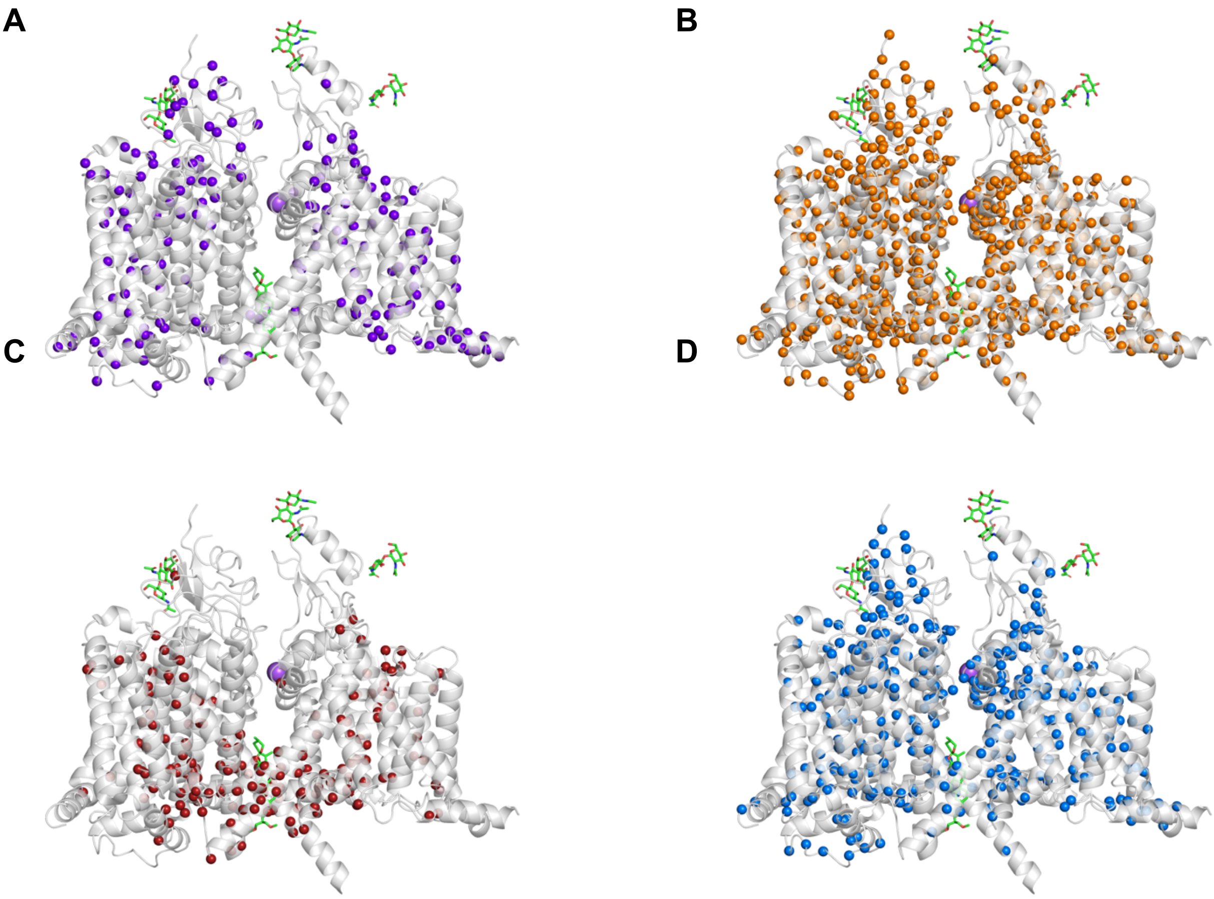

#### Figure S4, Related to Figure 3

**Quantitative protein features of GOF, LOF and neutral variants.** In this figure, we show histograms of quantitative protein features in the categories: accessible surface area, amino acid deleteriousness and physicochemical properties, conservation, secondary structure and variant density, separate for LOF, GOF and neutral variants. To account for different variant counts, we downsampled to the same numbers of LOF (blue), GOF (red) and neutral (purple) variants for SCN genes and CACNA1 genes, respectively. In total, we show 221 LOF, 221 GOF and 221 neutral SCN variants and 88 LOF, 88 GOF and 88 neutral CACNA1 variants. Variant density only includes variant density in *other* genes.

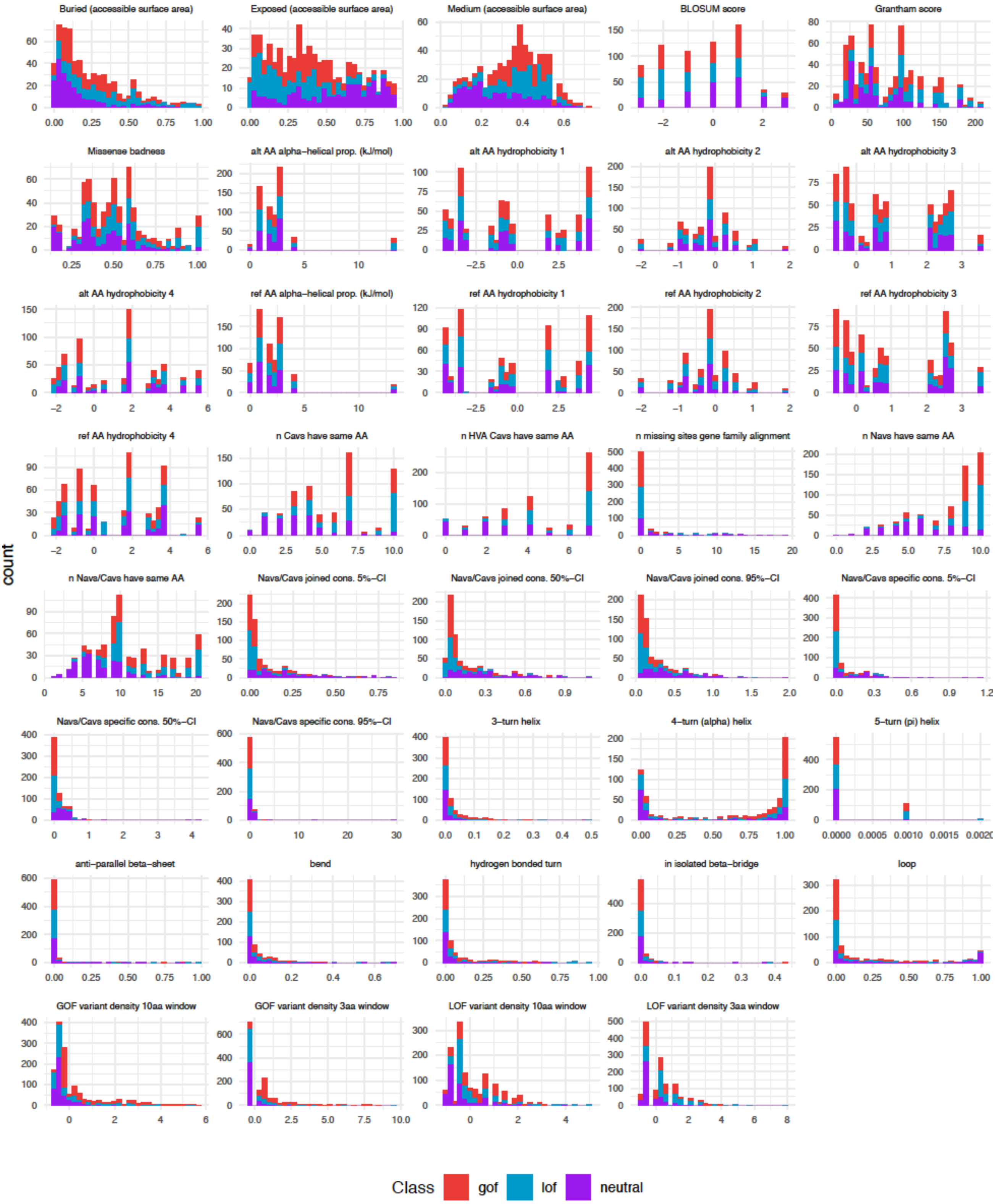
Buried (accessible surface area)
Exposed (accessible surface area)
Medium (accessible surface area)
BLOSUM score
Grantham score
Missense badness
alt AA alpha–helical prop. (kJ/mol)
alt AA hydrophobicity 1
alt AA hydrophobicity 2
alt AA hydrophobicity 3
alt AA hydrophobicity 4
ref AA alpha–helical prop. (kJ/mol)
ref AA hydrophobicity 1
ref AA hydrophobicity 2
ref AA hydrophobicity 3
ref AA hydrophobicity 4
n Cavs have same AA
n HVA Cavs have same AA
n missing sites gene family alignment
n Navs have same AA
n Navs/Cavs have same AA
Navs/Cavs joined cons. 5%–CI
Navs/Cavs joined cons. 50%–CI
Navs/Cavs joined cons. 95%–CI
Navs/Cavs specific cons. 5%–CI
Navs/Cavs specific cons. 50%–CI
Navs/Cavs specific cons. 95%–CI
3–turn helix
4–turn (alpha) helix
5–turn (pi) helix
anti–parallel beta–sheet
bend
hydrogen bonded turn
in isolated beta–bridge
loop
GOF variant density 10aa window
GOF variant density 3aa window
LOF variant density 10aa window
LOF variant density 3aa window
count
Class gof lof neutral

#### Figure S5, Related to Figure 4

**Performance of statistical modelling. A)** Performance (accuracy, kappa) comparison of different machine learning methods. **B)** Distribution of prediction accuracy during model training of the method “stochastic gradient boosting” **C)** Model performance (accuracy) in relationship to the tuning parameters “number of boosting iterations” (= trees) and “tree depth”. Performance was assessed with a 10-fold cross-validation approach.

**A)**

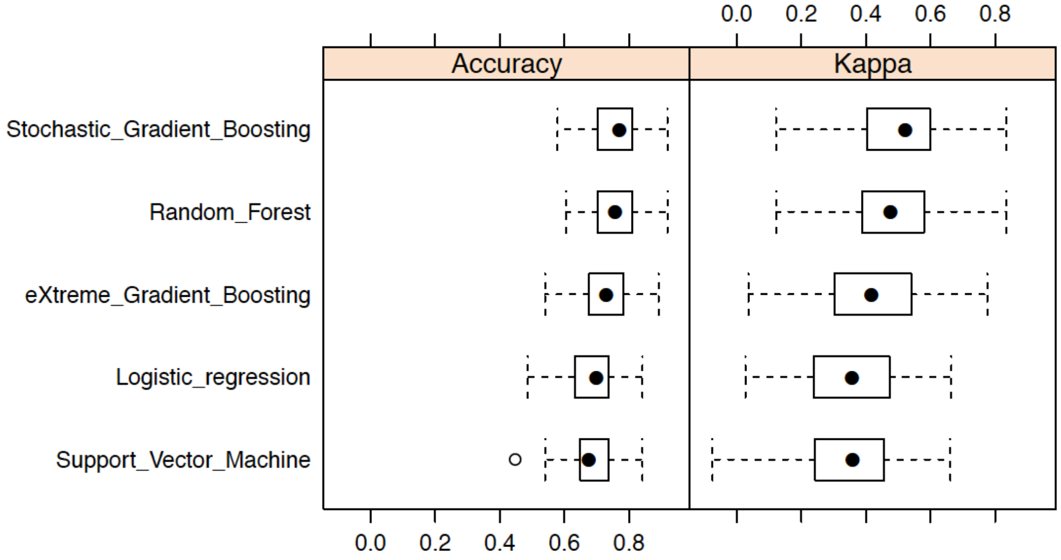


**B)**

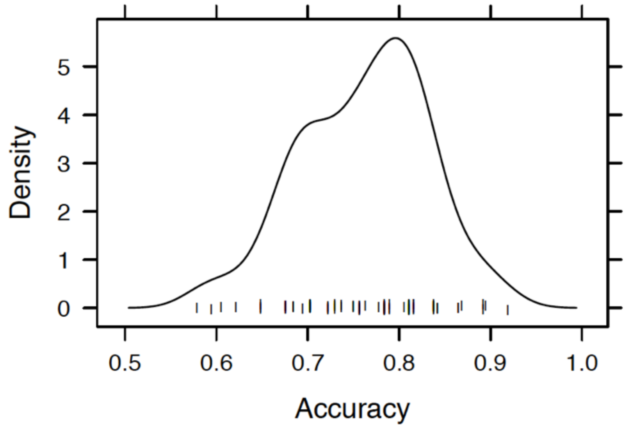


**C)**

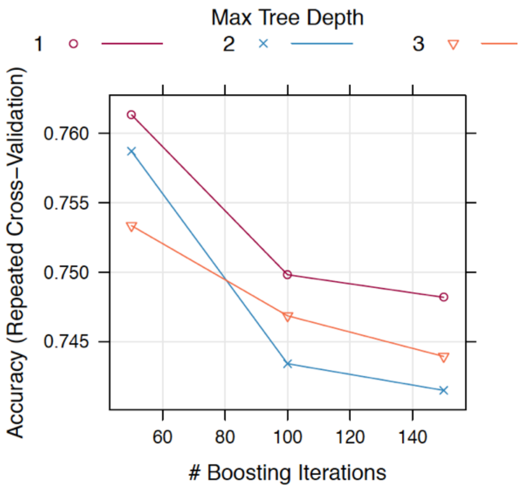

#### Figure S6, Related to Figure 5

**Prediction of pathogenic vs neutral variants.** Our statistical model (method GBM) trained on 4783 variants is used to predict an independent test data set of 530 variants. A) Receiver Operating Characteristic comparing prediction with our GBM model to three popular variant prediction tools MPC, PolyPhen-2 and CADD. The area under the Receiver Operating Characteristic (ROC) curve= 0.94. sensitivity=0.87, specificity=0.90. B) Feature importance plot. Relative influence of features on the prediction normalized to sum to 100 is computed as described in (Friedman, 2001). Of 89 features that went into the prediction, only the 37 features are shown that have a relative influence > 0.05% on the prediction.

**A)**

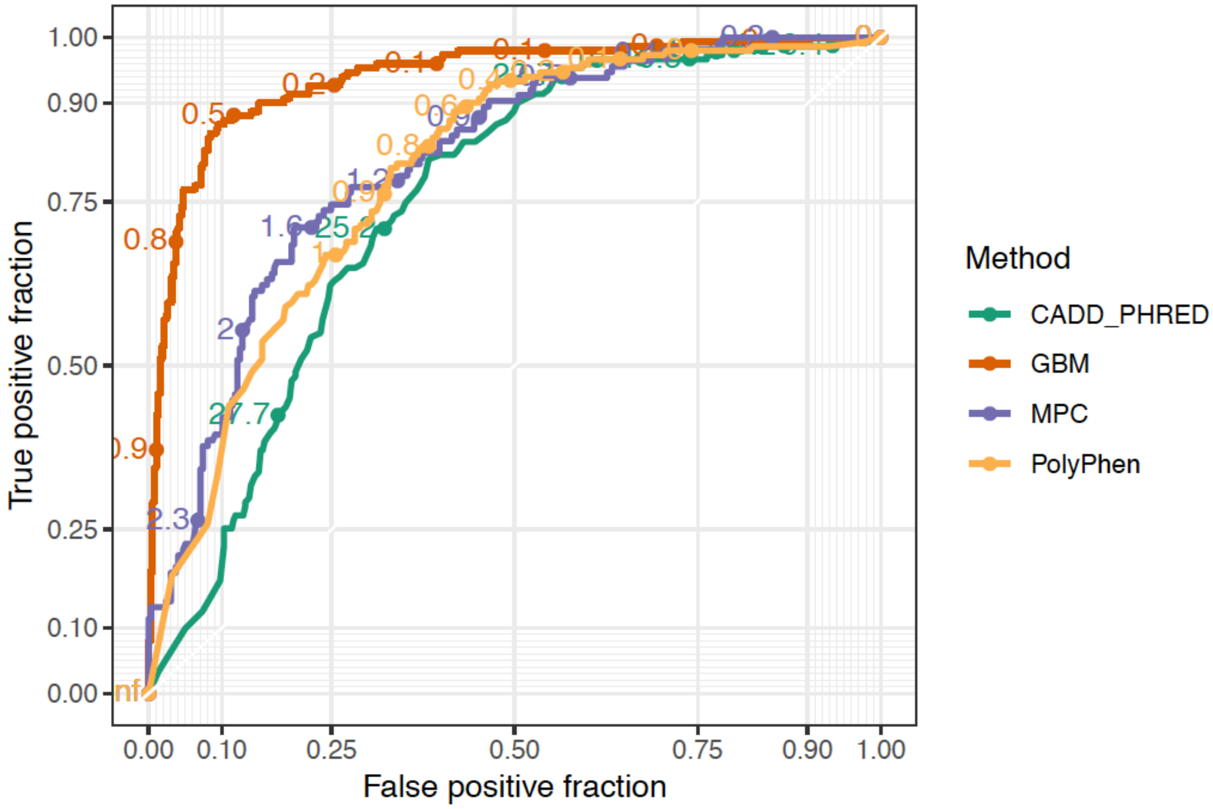

**B)**

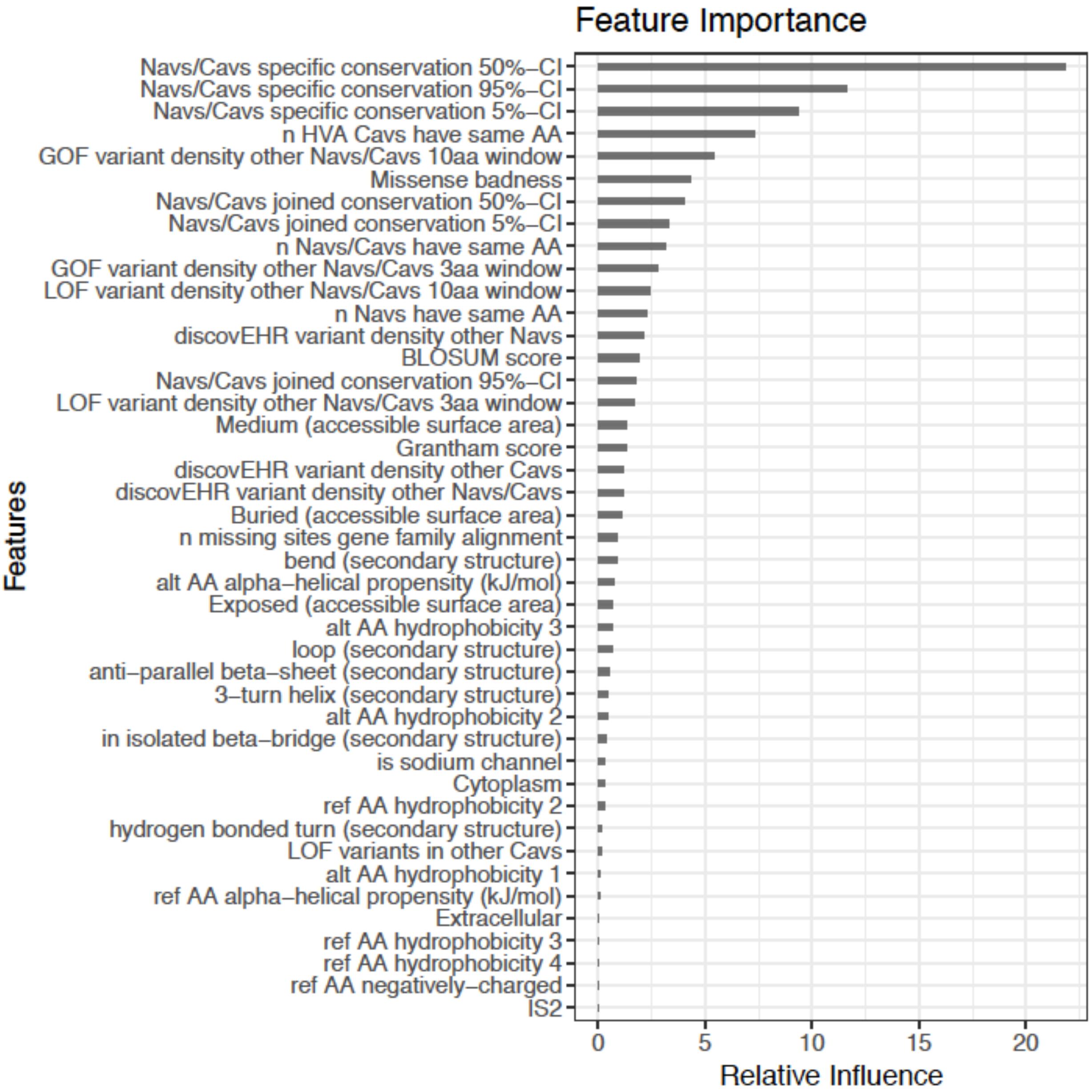
Feature Importance
Navs/Cavs specific conservation 50%–CI
Navs/Cavs specific conservation 95%–CI
Navs/Cavs specific conservation 5%–CI
n HVA Cavs have same AA
GOF variant density other Navs/Cavs 10aa window
Missense badness
Navs/Cavs joined conservation 50%–CI
Navs/Cavs joined conservation 5%–CI
n Navs/Cavs have same AA
GOF variant density other Navs/Cavs 3aa window
LOF variant density other Navs/Cavs 10aa window
n Navs have same AA
discovEHR variant density other Navs
BLOSUM score
Navs/Cavs joined conservation 95%–CI
LOF variant density other Navs/Cavs 3aa window
Medium (accessible surface area)
Grantham score
discovEHR variant density other Cavs
discovEHR variant density other Navs/Cavs
Buried (accessible surface area)
n missing sites gene family alignment
bend (secondary structure)
alt AA alpha–helical propensity (kJ/mol)
Exposed (accessible surface area)
alt AA hydrophobicity 3
loop (secondary structure)
anti–parallel beta–sheet (secondary structure)
3–turn helix (secondary structure)
alt AA hydrophobicity 2
in isolated beta–bridge (secondary structure)
is sodium channel
Cytoplasm
ref AA hydrophobicity 2
hydrogen bonded turn (secondary structure)
LOF variants in other Cavs
alt AA hydrophobicity 1
ref AA alpha–helical propensity (kJ/mol)
Extracellular
ref AA hydrophobicity 3
ref AA hydrophobicity 4
ref AA negatively–charged
IS2
Features
0
5
10
15
20
Relative Influence

#### Figure S7, Related to Figure 6

**Prediction of functionally tested variants**. The x-axis shows the GOF probability as predicted by our method. The horizontal grey line indicates the cutoff above which variants were labelled as GOF. Variants are colored by results of functional tests (electrophysiology). Of six duplicate variants in BFIS, two had opposite effects in different studies. Therefore we excluded variants in BFIS to benchmark our method.

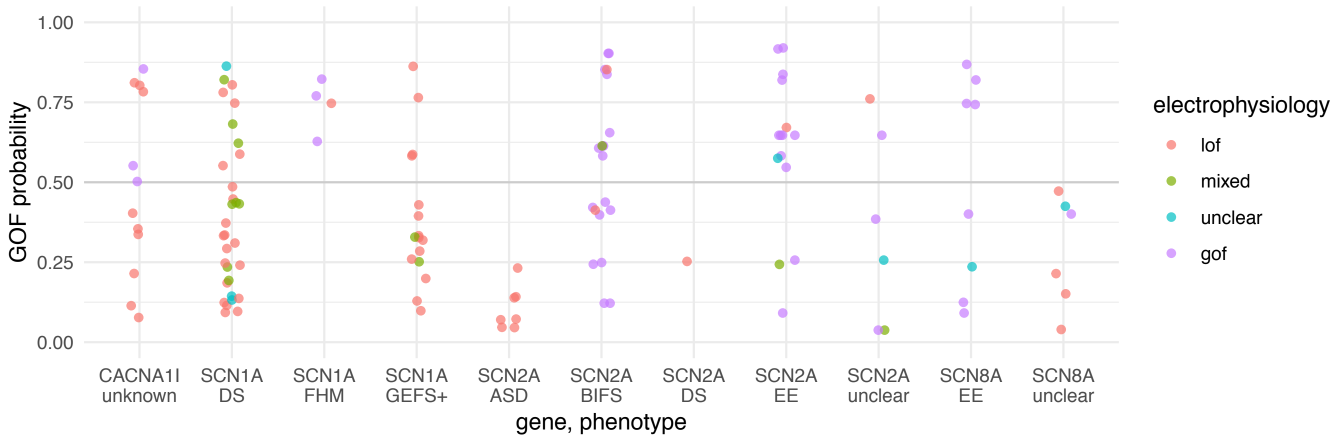

#### Figure S8

**Current Density Comparison of $Na_V$1.2 Wild-type and mutants.** $Na_V$1.2 K1422E current density is significantly lower than that of $Na_V$1.2 WT and $Na_V$1.2 K1420M. All experiments were done within the same day and quality controls were performed (seal resistance >500mohm, cell capacitance >2pF). Total numbers of cells tested are given as n in bars. Results of the experiment were replicated in two independent experiments (data not shown).

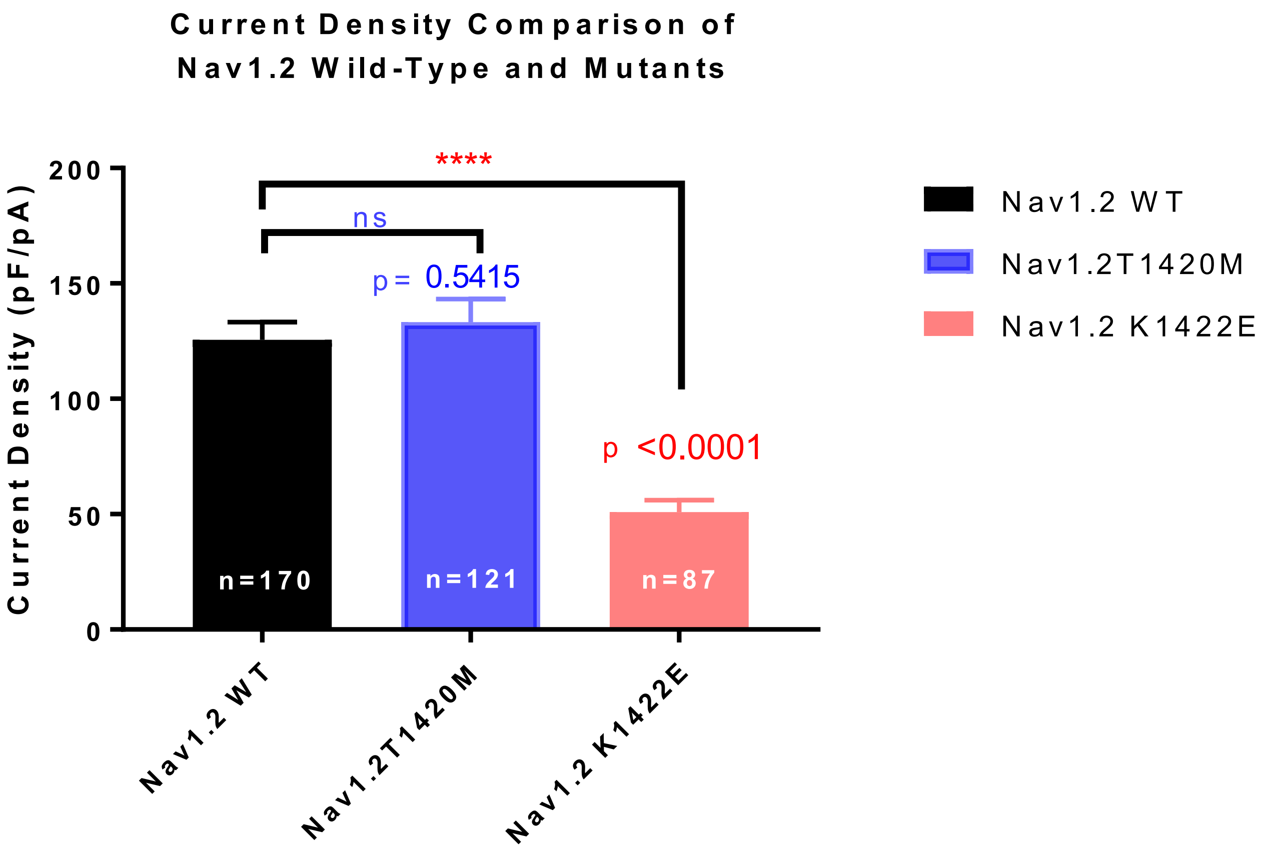
